# Toxic cocktails in soils - Evidence for synergistic effects of the imidacloprid-epoxiconazole mixture on earthworm life-history traits

**DOI:** 10.64898/2026.02.24.707680

**Authors:** Lisa Gollot, Cleo Tebby, Laura Frattaroli, Rémy Beaudouin, Raphaël Royauté, Juliette Faburé

## Abstract

Soils are vital reservoirs of biodiversity and providers of ecosystem services, yet they are increasingly threatened by agricultural intensification and pesticide use. Residues often persist as complex mixtures, while environmental risk assessment still largely focuses on single substances, potentially underestimating mixture effects. Earthworms play a key role in soil functioning and are particularly vulnerable to pesticide contamination. We investigated the effects of a binary mixture of epoxiconazole and imidacloprid, two persistent and frequently detected pesticides, on life-history traits of *Aporrectodea caliginosa*. We estimated each compound relative potency using dose–response experiments on juvenile growth and cocoon production. Next, we assessed the potential for synergy or antagonism in a fixed-ratio ray design including five concentration ratios and seven additive isoboles (36 conditions). Both compounds showed significant toxicity. Imidacloprid showed high potency (juvenile growth NOEC = 0.28 mg/kg; reproduction EC_50_ = 0.55 mg/kg), whereas epoxiconazole had moderate effects (juvenile growth NOEC = 9.3 mg/kg; reproduction EC_50_ = 126.8 mg/kg). Reproductive endpoints were more sensitive than adult growth, with juvenile growth being the most sensitive overall. Mixture analysis using Jonker’s models revealed significant deviation from Independent Action only under the simple interaction model, indicating synergism, consistent with cytochrome P450 interference reported in other taxa. Field-reported imidacloprid concentrations often approach effect thresholds, suggesting potential risks for earthworm populations. Overall, the combined effects of epoxiconazole and imidacloprid may exceed predictions not taking interactions into account. These results highlight the need to incorporate pesticide mixture effects into environmental risk assessment.

**Environmental Implications:** Pesticide residues persist in agricultural soils as complex mixtures, yet risk assessment still focuses mainly on single substances. This study shows that the combined effects of imidacloprid and epoxiconazole on earthworm reproduction can exceed predictions based on Independent Action, with evidence of synergistic interactions. Effect thresholds for imidacloprid approach reported maximum environmental concentrations, indicating limited safety margins for soil organisms. These findings suggest that mixture exposures may pose greater ecological risks than currently anticipated and highlight the need to integrate pesticide mixture toxicity and potential synergism into environmental risk assessment frameworks.

**Graphical abstract:** 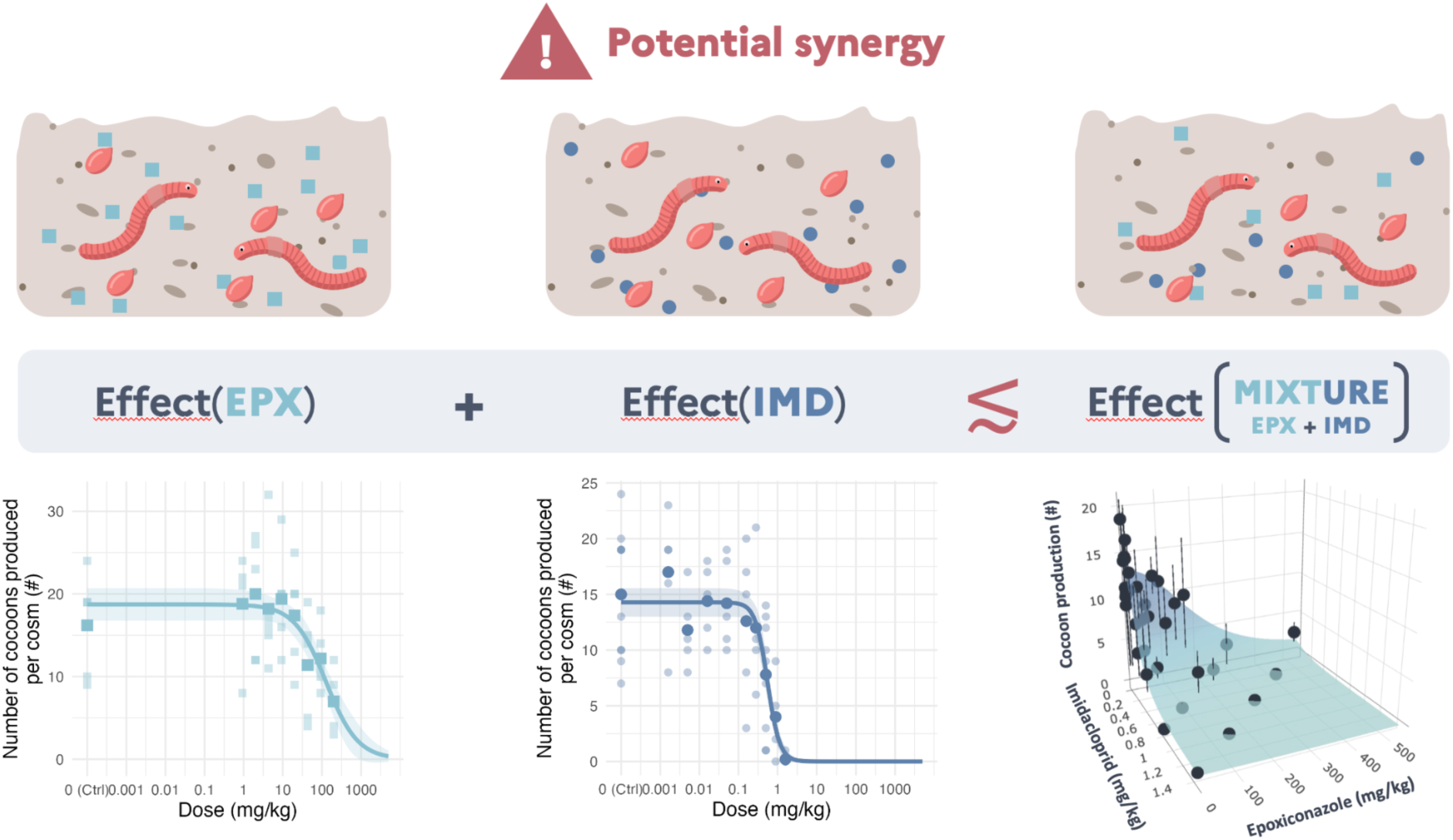

## 1. Introduction

Since the development of synthetic chemistry in the 19th century, pesticide use has steadily increased to support agricultural yields. In 2019, about 2.2 million tonnes of pesticides were used worldwide (Bondareva & Fedorova, 2021) and approximately 40,000 tonnes are used each year in France (MTE, 2022). This intensification has led to major contamination of agricultural soils: over 80% of tested soils in the European Union (EU) contain pesticides residues (Silva et al., 2019) and nearly all French agricultural soils may be contaminated by at least one pesticide (Pelosi et al., 2021). Soil-dwelling organisms are therefore highly exposed. As soil engineers and the dominant biomass in most terrestrial ecosystems (Lavelle et al., 2006), earthworms are particularly vulnerable to pesticides contamination since the soil represents both their habitat and food source. Pesticides have thus been identified as partly responsible for the decline in earthworm biodiversity in intensive agricultural areas (Bertrand et al., 2024; Smith et al., 2008).

Pesticides encompass a diverse array of chemical classes with distinct modes of action including neurotoxic organophosphates and carbamates that target acetylcholinesterase (Fukuto, 1990), neonicotinoids that act on nicotinic acetylcholine receptors (Tomizawa & Casida, 2003), pyrethroids that disrupt sodium channels (Soderlund & Knipple, 2003), azole fungicides that inhibit ergosterol biosynthesis (Price et al., 2015), and strobilurin that block mitochondrial respiration (Bartlett et al., 2002). In agricultural practice, most plant protection products consist of multiple active ingredients intentionally combined to broaden pest control efficacy and delay the development of resistance. This chemical and mechanistic diversity creates substantial potential for synergistic or antagonistic interactions when pesticides co-occur through processes affecting absorption, distribution, metabolism, or excretion or target site interactions (Cedergreen, 2014). Such interactions may emerge at multiple biological levels, but strong synergies often arise from metabolic interference, where one compound inhibits or enhances the biotransformation of another, altering persistence, activation, and ultimately toxicokinetics and toxicodynamics (Cedergreen, 2014).

Understanding the effects of chemical mixtures has therefore become a critical research priority. Two reference models are commonly used to predict the expected toxicity in the absence of interactions: Concentration Addition (CA) and Independent Action (IA). CA assumes that compounds act on the same biological target and contribute to the overall effect in proportion to their individual potencies (Greco et al., 1995). IA, in contrast, assumes that chemicals act through different modes of action and that their effects combine probabilistically. Under both frameworks, predicted mixture effects serve as a no-interaction baseline against which empirical responses are compared. Deviations from these expectations are classified as synergism (effects exceed predictions), or antagonism (effects fall below predictions). Potentiation represents a specific form of synergy in which a compound with little or no individual toxicity markedly increases the effect of another. Although non-interaction models provide strong predictive capacity for many environmental mixtures, this framework can fail when chemicals highly influence each other’s toxicokinetics or toxicodynamics, leading to unexpected synergistic or antagonistic effects (Cedergreen, 2014). Such interactions remain a major concern for ecological risk assessment because they challenge the assumption that mixture effects can be reliably inferred from single-compound data. To better characterize these deviations, more sophisticated modeling approaches have been developed, including the Jonker et al. (2005) simple interaction model, which provides a statistical framework for quantifying and characterizing deviations from the non-interaction models and therefore enabling robust interaction assessment.

Several studies (Froger et al., 2023; Pelosi et al., 2021) identified pesticides residues with the highest prevalence in French agricultural soils, including epoxiconazole (EPX) and imidacloprid (IMD), making them priority compounds for mixture toxicity assessment. Epoxiconazole is a broad-spectrum triazole fungicide that acts as a sterol biosynthesis inhibitor, while imidacloprid is a systemic neonicotinoid insecticide that targets nicotinic acetylcholine receptors, causing neurotoxicity through disrupted neural communication (Chen et al., 2021; Lewis et al., 2006). Although both compounds are now banned in the EU, their environmental persistence means they may continue to pose risks to soil organisms for years to come.

Both pesticides have demonstrated significant toxicity to earthworm species. Epoxiconazole exposure through formulations like Opus® and Swing Gold® causes substantial reductions in growth and reproductive output, with studies reporting up to 50% decreases in cocoon production at 3 times the former recommended dose, delayed hatching times, and reduced weight gains (Bart et al., 2019; Pelosi et al., 2016). At the biochemical level, epoxiconazole triggers oxidative stress and depletes energy reserves, indicating metabolic costs associated with detoxification processes (Givaudan et al., 2014; Pelosi et al., 2016). Similarly, imidacloprid impairs earthworm reproduction and growth across multiple species, with effects on cast production showing concentration-dependent responses—low concentrations (0.2 mg/kg) initially increase activity, while higher concentrations (≥ 0.66 mg/kg) cause dramatic declines (Dittbrenner et al., 2010; Van Loon et al., 2022; Wang et al., 2019). At the infra-individual level, imidacloprid interferes with multiple physiological pathways in earthworms. It impairs detoxification by inhibiting carboxylesterase, leading to reduced detoxification capacity (Wang et al., 2019). At the cellular level, it induces genotoxic effects such as DNA damage and sperm deformities in *Eisenia fetida* (Kaur et al., 2023; Zang et al., 2000). Together, these findings suggest that imidacloprid compromises detoxification, reproduction-related neuroendocrine signaling, and genome integrity in earthworms.

Beyond their individual toxicological profiles, these compounds could interact in ways that modulate their combined toxicity. Triazole fungicides such as epoxiconazole are well-documented inhibitors of cytochrome P450 monooxygenases, key enzymes involved in xenobiotic metabolism (Bass et al., 2015; Cedergreen et al., 2006; Puinean et al., 2010). By inhibiting these enzymes, triazoles can slow biotransformation and elimination of co-occurring insecticides, maintaining elevated internal concentrations and amplifying toxic effects. In *Daphnia magna*, such metabolic inhibition enhanced esfenvalerate toxicity by up to fourfold (Cedergreen et al., 2006) and increased α-cypermethrin toxicity threefold (Gottardi et al., 2017). Synergistic effects between prochloraz and deltamethrin, as well as between several other sterol biosynthesis-inhibiting fungicides and thiacloprid, have also been observed in the honeybee (*Apis mellifera*) (Meled et al., 1998 ; Schmuck et al., 2003). Although imidacloprid is moderately hydrophilic (log K_ow_ ≈ 0.57), evidence indicates partial metabolism via cytochrome P450 enzymes in several insect species (Chen et al., 2018; Chen et al., 2019). Cytochrome P450 overexpression has been linked to imidacloprid resistance in multiple insect species (Elzaki et al., 2017; Kaplanoglu et al., 2017; Liang et al., 2015), suggesting that comparable metabolic pathways may exist in other invertebrates like earthworms. These findings suggest that comparable interactions between epoxiconazole and imidacloprid could occur in earthworms, potentially resulting in synergistic effects on growth, reproduction, and survival through elevated internal exposure to the insecticide.

To investigate these potential mixture interactions, we selected *Aporrectodea caliginosa* as our model species. This widely distributed endogeic earthworm plays a crucial role in soil ecosystem functioning through its burrowing activity and organic matter processing. The species is commonly found in agricultural soils across Europe and represents a relevant model organism for ecotoxicological studies due to its ecological importance and demonstrated sensitivity to pesticides contamination (Bart et al., 2018; Pelosi et al., 2013).

The objective of this study was to characterize the individual and combined toxicity of epoxiconazole and imidacloprid to *A. caliginosa* life-history traits using standardized reproduction and growth bioassays. We employed a ray-design experimental approach to systematically assess mixture interactions across different effect-based ratios, followed by statistical modeling to quantify deviations from additivity and from independence of action and identify the nature of any interactions present.

Given the widespread occurrence of epoxiconazole and imidacloprid in agricultural soils and the potential for metabolic interactions between triazole fungicides and neonicotinoid insecticides, several research questions emerge. First, how do the dose-response relationships of each of these compounds compare in terms of toxicity to *A. caliginosa*? Second, when present as a binary mixture, do epoxiconazole and imidacloprid exhibit mixture effects as predicted by models without interactions, or do they interact synergistically or antagonistically? Finally, can these mixture interactions be adequately characterized using existing modeling frameworks?

We hypothesize that epoxiconazole and imidacloprid will exhibit different individual toxicity profiles due to their distinct modes of action, with imidacloprid showing greater potency based on its neurotoxic mechanism. Furthermore, we hypothesize that the cytochrome P450 inhibition caused by epoxiconazole may enhance imidacloprid toxicity, leading to synergistic interactions that deviate from additivity or independent action predictions.

## 2. Materials and methods

To evaluate potential interactions between the two pesticides, we applied a sequential experimental design integrating single-compound dose–response characterization and mixture testing, associated with response-surface modeling. First, individual dose–response curves were modelled to derive key ecotoxicological parameters. These informed a ray design mixture experiment spanning 5 fixed effect ratios (100:0 to 0:100). Finally, the combined effects were analyzed using the interaction models proposed by Jonker et al. (2005), which compares observed responses with concentration addition and independent action predictions to detect synergistic or antagonistic deviations.

### 2.1 Experimental material

Laboratory-bred earthworms originated from individuals collected in late 2023–early 2024 from a natural population in a permanent grassland in Versailles (48°48′ N, 2°5′ E). No pesticides have been applied at this site for more than 30 years. Individuals were identified using a binocular microscope and an identification key. Individuals were bred at 18°C in 1 L vessels in groups of five individuals using a soil sampled from this same meadow, a loamy soil texture (based on the texture definition of the Food and Agriculture Organization of the United Nations (FAO)), to be as close as possible to the conditions of the natural population (see Table S1.1 for soil characteristics). The soil was collected from the top 0-20 cm, air-dried and crushed to pass a 2 mm mesh. Soil culture was kept with a soil water-holding capacity (WHC) of 60-70% and individuals were fed with horse dung collected in-between veterinary deworming treatments and then frosted and defrosted twice before being milled (< 1 mm; Lowe & Butt 2005). The equivalent of 6 g of dry horse dung per group of five earthworms were given with a WHC of 60-80% every month. As recommended by Bart et al. (2018), ten individuals from the natural population were barcoded, showing individuals come from two lineages of the *Aporrectodea caliginosa* species (see Figure S1.1). Earthworms used in each reproduction test were less than two months apart in age, as reproductive output varies with age (Hartenstein et al., 1979).

For all experiments, Imidacloprid PESTANAL®, analytical standard (purity: ≥ 98.0%, ref 37894-100MG) and Epoxiconazole, analytical standard (purity: 98.75%, ref HY-119683) from Sigma-Aldrich were used. Due to the very low water solubility of epoxiconazole and the relatively high tested concentrations, it was not possible to spike the soil with a water solution containing both epoxiconazole and imidacloprid. The alternative use of a co-solvent was avoided, as they are often toxic to earthworms, and the required quantity would have been excessive. One day prior to the start of the tests, the soil contamination was carried out in two stages: first, a dry mixture of epoxiconazole and soil at 20% of its WHC was prepared, followed by spiking the soil with a water solution of imidacloprid. Soil samples were collected at the start of each experiment for every prepared soil batch to determine the actual exposure concentration.

### 2.2. Earthworms toxicity tests

Each pesticide was first tested individually (UNI experiment) before assessing the mixture (MIX experiment, see section 2.3).

#### 2.2.1 Reproduction test

Reproduction tests were conducted following an adaptation of the OECD Technical Guideline 222 (OECD, 2016) in a two-stage experiment.

For each replicate, two adult earthworms were placed in a cosm consisting of a plastic vessel (11 cm × 7.7 cm × 4.5 cm) containing the equivalent of 200 g of dry soil. The exposure lasted 28 days at 18°C. Earthworms were weighed at the start and end of the experiment and were fed ad libitum with horse dung (3 g per individual every 14 days). Soil moisture was checked and adjusted weekly. Cocoons produced during the experiments were retrieved by wet-sieving the soil through a 1-mm mesh (Bart et al., 2018) and incubated at 18°C in Petri dishes on wet filter paper (Holmstrup et al., 1991). Cocoon hatching was monitored each day of the working week for 8 weeks.

#### 2.2.2. Juvenile growth test

Juveniles of similar body weights were sampled from the laboratory culture (mean = 97.0 mg ; sd = 35.6 mg). Juvenile growth tests followed the same pattern as reproduction tests in the same conditions of temperature, soil moisture and food supply. For each replicate, one earthworm was placed in a cosm consisting of a plastic vessel (11 cm × 7.7 cm × 4.5 cm) containing the equivalent of 200 g of dry soil. Growth rates of each individual were estimated using a linear regression on the structural length of the earthworm 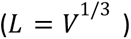 which is related to the weight without gut content (*W*) through an allometric relationship with a 1/3 power scaling (Gollot et al., 2025).

### 2.3. Single substances testing

The first step of our study involved evaluating each substance individually on each endpoint, as characterizing single-substance effects is a prerequisite for understanding mixture interactions. We assessed both reproductive endpoints through cocoon production in adult earthworms and developmental endpoints through juvenile growth rates, providing a first evaluation of each compound’s toxicological profile before proceeding to mixture testing (UNI experiment).

To study the effects of individual pesticides, tested concentrations were selected based on the ecotoxicological data from Lewis et al. (2006). For *Eisenia fetida*, the reported acute 14-day LC_50_ ranged from 10.7 mg/kg for imidacloprid to >500 mg/kg for epoxiconazole, while the chronic reproduction NOEC values were≥ 0.178 mg/kg and ≥ 3.24 mg/kg respectively. An initial set of eight concentrations was selected for each pesticide, arranged in a geometric sequence to cover the full response range (Table 1). For imidacloprid, an additional set of two concentrations (C6 and C8) with an additional concentration already tested in the initial set (C7), was added to refine the dose-response curve. Each concentration was tested in five replicates.

**Table 1.** Tested concentrations of epoxiconazole and imidacloprid during the UNI experiment.

|  | C1 | C2 | C3 | C4 | C5 | C6 | C7 | C8 | C9 | C10 |
| --- | --- | --- | --- | --- | --- | --- | --- | --- | --- | --- |
| IMD<br>(mg/kg) | 0.0016 | 0.0050 | 0.016 | 0.050 | 0.16 | 0.28 | 0.50 | 0.89 | 1.6 | 5.0 |
| EPX<br>(mg/kg) | 0.9 | 2.0 | 4.3 | 9.3 | 20.0 | - | 43.1 | - | 92.8 | 200 |

Dose-response curves were modeled using the Hill log-logistic function. Growth rate and cocoon production dose-response curves were modeled using Normal and Poisson error distributions, respectively. The model fit of all dose-response models was evaluated using lack-of-fit tests compared to Anova models.

### 2.4. Substances in combination

#### 2.4.1. Modelling approach for mixture interaction assessment

Mixture effects were evaluated using two reference models: concentration addition (CA) and independent action (IA) also known as response addition, both of which assume that the effects of two substances are predictable from their individual dose–response functions without any interaction. These models do not introduce additional parameters and serve as the null hypothesis. CA assumes that chemicals behave as dilutions of one another and is most appropriate when compounds share a similar mode of action. CA is generally considered conservative, often predicting stronger mixture effects even when modes of action differ (Drescher & Boedeker, 1995). IA assumes that chemicals act through independent modes of action. It predicts the probability of effect based on the individual dose–response functions and is typically less conservative than CA. Using both CA and IA provides a complementary, widely used framework to detect potential synergistic or antagonistic deviations from additivity (see section S3 for more detail). The use of IA is warranted given the mechanistic dissimilarity between the two compounds: imidacloprid functions as a nicotinic acetylcholine receptor agonist, while epoxiconazole disrupts sterol biosynthesis.

If the CA model or the IA model adequately describe the mixture data, no interaction is inferred. However, when deviations occur, the framework introduces three interaction models that extend CA and IA with one or two additional parameters (see section S3 for details). In the *simple interaction* model (SA), a single parameter *a*, quantifies a constant shift from the CA or IA prediction, regardless of the dose level or mixture ratio. Here, *a* > 0 indicates a systematic synergistic deviation, *a* < 0 indicates antagonism, and *a* = 0 corresponds to strict additivity (the CA model). In the *dose-level dependent* model (DL), the parameter *b* modifies the predicted effect depending on the magnitude of the response. This allows interactions to strengthen or weaken at higher effect levels, capturing scenarios where, for example, synergy emerges only at high concentrations. In the dose-*ratio dependent model* (DR), the parameter *b* links the strength of the interaction to the relative proportions of the two compounds. This formulation captures cases where mixture toxicity depends strongly on whether one compound dominates the mixture composition.

#### 2.4.2. Experimental design

To enable proper fitting of the response-surface models, we designed our mixture experiment (MIX experiment) using a ray-design approach (Gessner et al., 1995; Greco et al., 1995) (Figure 2), which involves testing predefined mixture ratios based on toxic units derived from individual compound potencies. This experimental design ensures adequate coverage of the dose-response surface while providing the necessary data structure for reliable model calibration and interaction assessment. Although this design is built to ensure optimal detection and visualisation of deviations from concentration additivity, it also allows detection of deviations from the independence of action model.

This design assesses binary mixtures at predefined ratios based on Toxic Units:

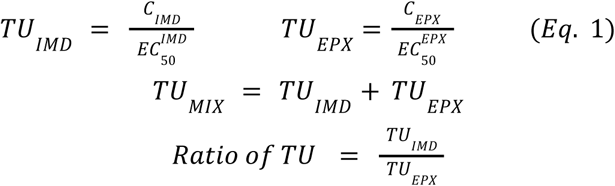

Where C_IMD_ is the imidacloprid concentration, C_EPX_ the epoxiconazole concentration, 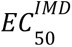 the effective concentration 50 of imidacloprid, 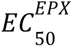 the 50% effective concentration of epoxiconazole and *TU_IMD_* and *TU_EPX_* the corresponding toxic unit of imidacloprid and epoxiconazole at given concentrations of imidacloprid and epoxiconazole.

Ratios of Toxic Units tested were: 1:0, 3:1, 1:1, 1:3, and 0:1. For example, a ratio of 3:1 at TU_mix_ = 1 means that the mixture corresponds to 0.75 TU of imidacloprid and 0.25 TU of epoxiconazole. In other words, mixture ratios were selected to distribute the contribution to the joint effect between the two pesticides in fixed proportions, from single-compound exposures (1:0 and 0:1) to equitoxic (1:1) and non-equitoxic combinations (3:1, 1:3). This design allows evaluation of whether deviations from concentration addition occur across the whole gradient of relative contributions. The concentrations of each pesticide required to achieve these effect-based mixture ratios were calculated from EC_50_ values derived from individual dose-response curves fitted to the UNI experiment data.

The individual dose-response curves derived from the UNI experiment data were constrained to have common baseline and maximal response levels (Y_min_, Y_max_), as well as equal slope parameters for both pesticides, as assumed in the Jonker et al. (2005) interaction models. The suitability of the three equal dose-response parameters for both pesticides was tested using likelihood ratio tests comparing models with compound-specific parameters against models with constrained equal parameters. The EC_50_ values from these constrained dose-response curves were then used to design the MIX experiment, ensuring that tested concentrations would theoretically match effect isoboles under the concentration addition (CA) hypothesis.

Because we were not able to produce complete dose-response curves for juvenile growth with the UNI experiment (see section 3.1.1.), mixture interaction assessment was only performed on cocoon production from reproduction tests. For the MIX experiment design, seven theoretical effect levels (effect isoboles) spanning the full response range were selected and calculated under the CA hypothesis (more detail available section S3.1.1.). These effect isoboles are defined by Equation 2.

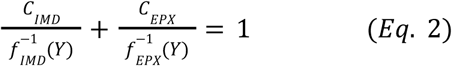

With Y the observed effect for an exposition to the mixture with a concentration *C*_*IMD*_ of imidacloprid and a concentration *C*_*EPX*_ of epoxiconazole, and 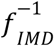 and 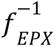 the inverse function of the dose-response relationship.

Including the five mixture ratios and the controls, a total of 36 experimental conditions were assessed, with each condition replicated four times to ensure statistical robustness and account for biological variability in earthworm responses.

For logistical reasons, the MIX experiment was conducted in two batches, with each batch representing two replicates of all treatment conditions using earthworms randomly selected from the same culture pool. The two batches were staggered by only one week. The reproduction tests followed the same protocol as the UNI experiment but with individual weight measurements facilitated by tagging earthworms with Visible Implant Elastomers (VIE, Northwest Marine Technology Inc.) two weeks prior to the experiment using the method described by Butt & Lowe, (2007). Cocoon hatching was monitored daily during working days for 8 weeks for the first three replicates to assess reproductive success beyond cocoon production.

#### 2.4.3. Mixture data and statistical analysis

To minimize the confounding influence of both mixture interactions and inter-experiment variability, the MIX experiment included single-substance concentration rays, following established recommendations (Dawson et al., 2017; Kappenberg et al., 2025). These rays were generated under the same experimental conditions as the mixture treatments, ensuring that single-compound dose–response data were fully comparable with mixture data.

Dose-response curves were fitted to single exposure data from the MIX experiment in order to check the underlying assumptions of an identical slope and identical minimum responses (*Y*_*min*_) and responses in control conditions (*Y*_*max*_) for both substances. This assumption was tested with a likelihood ratio test. Dose-response modeling of mixture data was only performed when these assumptions were satisfied and model fit was evaluated using lack-of-fit tests.

In practice, the procedure proposed by Jonker et al. (2005) to analyse mixture data consisted first in fitting the CA and the IA model under the assumption of common slope, *Y*_*min*_ and *Y*_*max*_ across dose-response curves. As the endpoint is cocoon production, a Poisson error model was used which implied reformulation of the loss function used during parameter optimization. Due to the structure of the error model, model comparison using F-tests was not possible. Instead, likelihood-ratio tests were employed to compare models. The adequacy of the CA and IA model was evaluated by comparing them to an Generalized Linear model with Poisson error - this ANOVA modelled the response as a function of the tested conditions using a Likelihood ratio test (lack-of-fit test). The CA and the IA model were then extended stepwise with interaction terms (*a* and optionally *b*), leading to the SA, DL, or DR formulations. The performance of these models was then compared against the CA or IA baseline using Likelihood ratio tests, in order to determine whether including interaction terms significantly improved model fit. Uncertainty around the interaction parameters (*a*, *b*) was quantified by bootstrap resampling (*n* = 5000 with the same number of draws per condition), which provided confidence intervals and allowed us to classify interactions as synergistic, antagonistic, or additive (section S3).

All data analyses were performed using R (R Core Team, 2015), with dose-response curves modeled using the ‘drc’ package (Ritz et al., 2015). The Jonker models were implemented using custom R code, as no existing package currently provides a direct implementation. The surface modelled by the Jonker model can be represented in 3D and also in 2D with its isoboles.

## 3. Results

### 3.1. Study of the single pesticides

#### 3.1.1. Effect on juvenile growth

Juvenile growth tests revealed substantial differences in sensitivity between the two compounds (Figure 1). Both pesticide exposures resulted in declining growth rates with increasing concentrations (F-tests, p-values < 1e-9 for both substances). Growth rates decreased by approximately 130% and 50% for IMD and EPX at the highest tested concentrations, respectively. The three highest tested concentrations of imidacloprid caused partial mortality, with 4/110 earthworms dying by the end of the experiment under these conditions. These earthworms were excluded from the analysis. The surviving earthworms exhibited negative growth (shrinking). In contrast, epoxiconazole showed more moderate effects on earthworm growth with no observed shrinking or mortality. For imidacloprid, we observed maximal growth inhibition at concentrations where growth effects were confounded by lethal toxicity. However, epoxiconazole effects remained moderate across the tested concentration range, suggesting that stronger effects might occur at higher doses. Higher concentrations were not tested due to the associated costs and their limited environmental relevance. Due to the lethal effects observed with imidacloprid at high concentrations and the failure to reach maximum effect levels with epoxiconazole, reliable dose-response curves could not be established for this endpoint. However, NOEC of 0.28 mg/kg and 9.28 mg/kg and LOEC of 0.5 and 20 mg/kg were determined for imidacloprid and epoxiconazole, respectively (Table 3).

**Figure 1.**
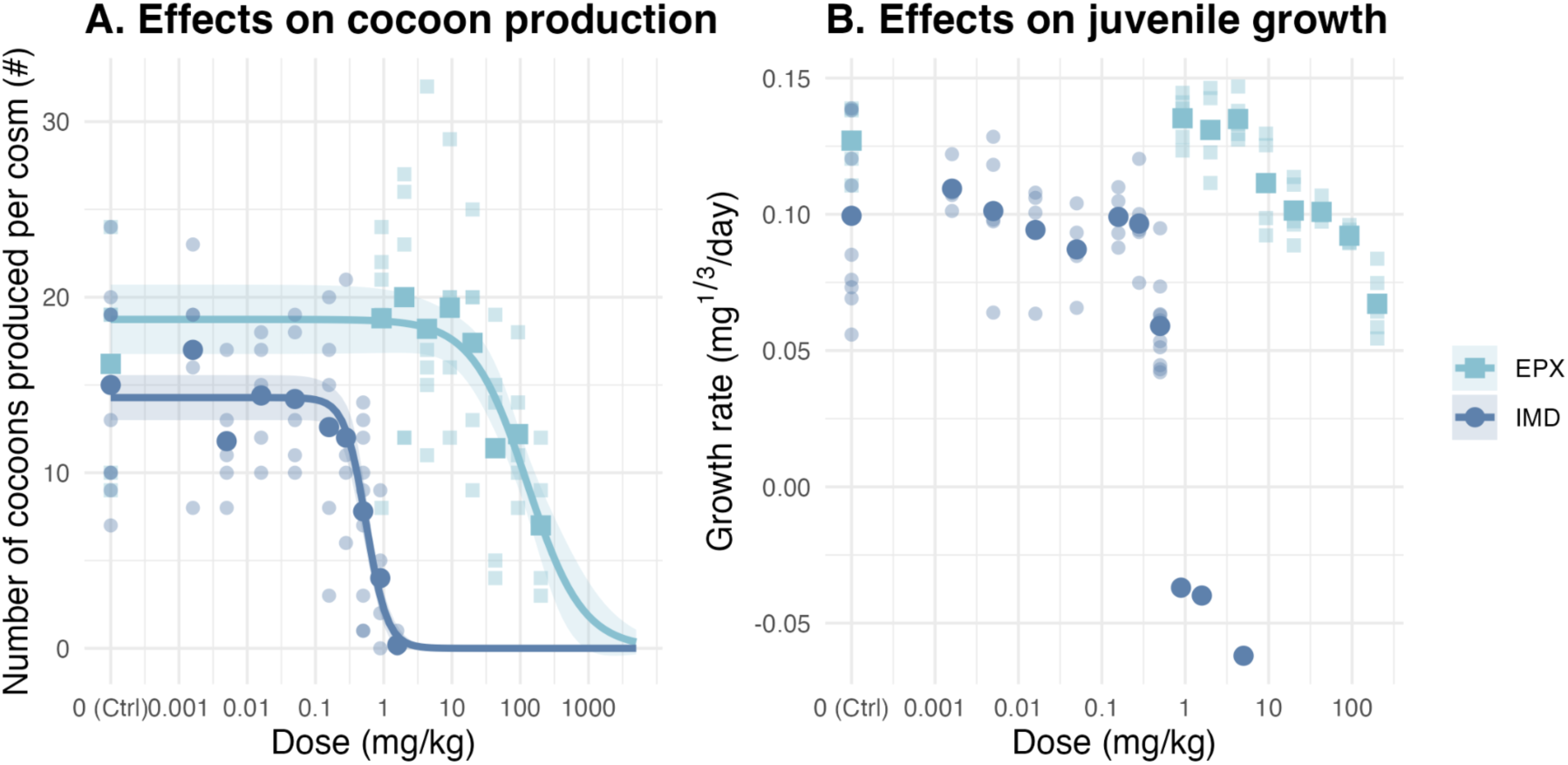
Effect of imidacloprid (dark blue line) and epoxiconazole (light blue line) obtained on earthworm cocoon production (panel A) and on juvenile growth (panel B). Experimental data points from the UNI experiment are represented by the transparent colored points while bolder points correspond to the mean per condition. Thick lines and their respective ribbons represent the estimated dose-response curves with their 95% confidence intervals.

#### 3.1.2. Effect on earthworm reproduction

Dose-response curves for the cocoon production endpoint were successfully established for the two compounds (Figure 1). From these curves, we derived EC_50_ and NOEC values that revealed again substantial differences in sensitivity between the two compounds (Table 3). The EC_50_ for imidacloprid was 0.55 mg/kg (CI 95% : [0.45 ; 0.65]), while the EC_50_ for epoxiconazole was 126.8 mg/kg (CI 95% : [73.6 ; 180]), indicating that earthworms are considerably more sensitive to imidacloprid than to epoxiconazole (Table S2.2). The EC_50_ ratio (EPX/IMD) indicates imidacloprid is 230 times more potent than epoxiconazole in reducing cocoon production. Under control conditions, earthworms produced an average of 13.8 and 16.2 cocoons per cosm across the two experimental batches, respectively, establishing baseline reproductive output for comparison with treated groups to derive NOEC and LOEC (Table 3). We could additionally determine NOEC for the adult growth endpoint with values of 0.5 mg/kg and 92.8 mg/kg for IMD and EPX, respectively.

### 3.2. Effects of pesticides interaction

#### 3.2.1. Determination of the experimental design

As dose-response relationships for juvenile growth could not be established, mixture experiments could not be conducted for this endpoint. This limitation arose because the maximum effect could not be properly defined, in contrast to reproduction, where the absence of cocoon production clearly represents the maximal effect.

Mixture experiments monitoring cocoon production were designed by first obtaining reliable EC₅₀ estimates for each compound individually, then deriving the concentrations required to achieve the effect-based mixture ratios. To ensure compatibility with the dose-response surface modelling workflow described by Jonker et al. (2005), the compound-specific dose–response curves were re-fitted with common *Y*_*min*’_, *Y*_*max*’_ and *Slope* parameters for both pesticides. Likelihood-ratio tests indicated that the model fit of these constrained models did not differ significantly from the fully compound-specific ones, thereby validating the use of identical parameter values. This allowed us to establish a robust experimental design for experiment MIX (Figure 2).

**Figure 2.**
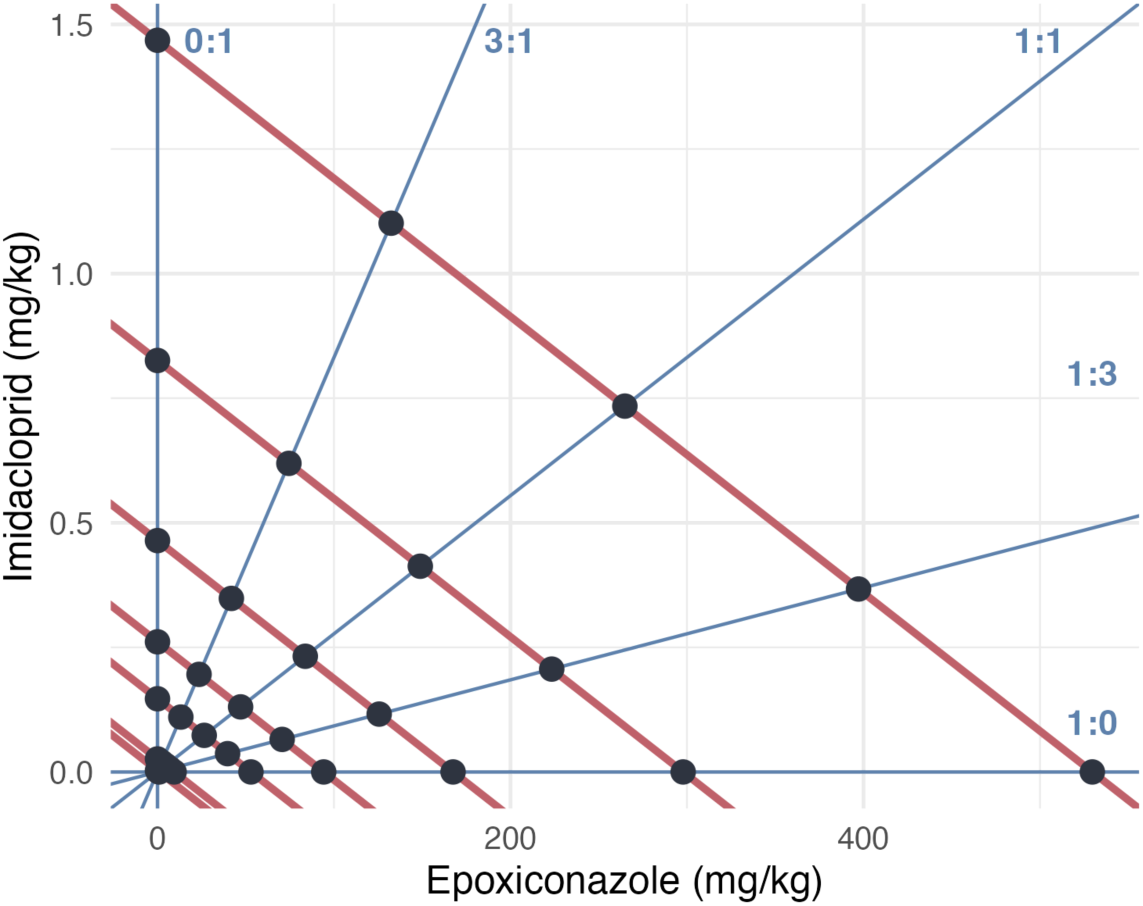
Experimental design for the binary mixture of imidacloprid and epoxiconazole. Black points correspond to the tested pairs of concentrations. Blue lines represent the effect-based mixture ratios and the red lines represent the theoretical effect isobole, assuming the concentration addition hypothesis is valid.

Power analysis revealed clear thresholds in the ability of the simple interaction (SA) model to detect and quantify interactions. With CA as the reference model, interaction strengths of |*a*| ≈ 1 (see Section S3.1 for interpretation of parameter *a*) provided adequate power (∼97%) and acceptable estimation accuracy (< 20% error), whereas performance degraded markedly at |*a*| = 0.5. Using IA as the reference yielded the same qualitative threshold: |*a*| ≈ 1 was the minimum interaction strength at which detection remained reasonably reliable, although power was lower (∼80%) than under CA. For both reference models, weaker interactions (|*a*| ≤ 0.5) led to insufficient power and poor estimation accuracy, making them difficult to detect or classify robustly. Full numerical details are provided in Table 2 and Section S3.3.

**Table 2.** Estimated interaction parameter for each mixture model and their respective negative log-likelihood.

| Model | $a$ | $b$ | Log-likelihood | p-value<br>(Likelihood-ratio test<br>against reference<br>model) |
| --- | --- | --- | --- | --- |
| CA | - | - | -395.2 | - |
| CA-SA | -0.005 [-1.06 ; 0.84] | - | -395.2 | 0.992 |
| CA-DL | 1.6 [-0.7 ; 5.5] | 0.63 [-174.4 ; 0.65] | -392.3 | 0.052 |
| CA-DR | -0.40 [-3.4 ; 2.3] | 0.9 [-5.1 ; 6.7] | -395.1 | 0.892 |
| IA | - | - | -397.5 | - |
| IA-SA | -1.5 [-3.0 ; -0.2] | - | -391.3 | 0.0004* |
| IA-DL | -1.9 [-6.8 ; -0.004] | 0.6 [-298.5 ; 2.3] | -391.1 | 0.002*<br>Against IA-SA :<br>0.52 |
| IA-DR | -1.6 [-5.6 ; 2.1] | 0.3 [-8.2 ; 8.1] | -391.3 | 0.002*<br>Against IA-SA :<br>0.91 |

#### 3.2.2. Mixture interaction analysis

Results of the MIX experiment were represented as dose-response curves for each mixture ratio with the number of cocoons plotted against the TU of the mixture (Figure 3). Under the assumptions of equal slopes, maximum and minimum response levels and concentration additivity, the curves should be identical.

**Figure 3.**
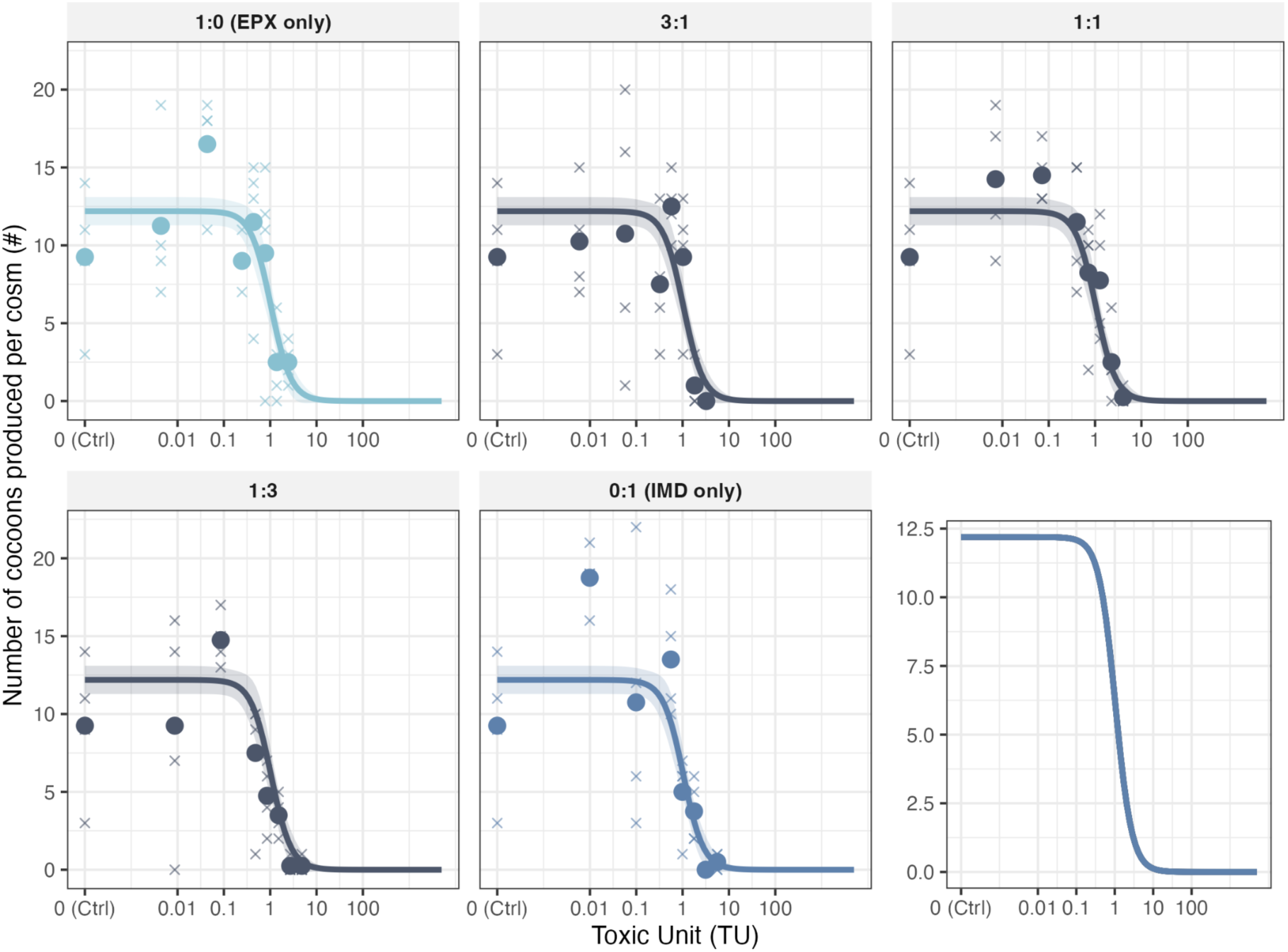
Dose-response curves obtained for each mixture ratio. The lines represent the dose-response curves obtained (with common *Y*_max_ and *Y*_min_ set to 0). Crosses correspond to the number of cocoons produced by one cosm and the bold points represent the mean per condition. The bottom-right graphic shows the superposition of the dose-response curves for all ratios.

The CA and IA reference models were successfully determined (Figure 4). Analysis of the mixture dose-response data with the methodology proposed by Jonker et al. (2005) revealed no significant deviations from the CA model. In other words, the Likelihood-ratio tests between CA model and interaction models were not significant. The CA-SA model yielded an interaction parameter of a = −0.005 [−1.08 ; 0.84], indicating a non significant deviation from additivity. Likelihood-ratio tests against the concentration addition null hypothesis showed no statistical significance (Table 2).

**Figure 4.**
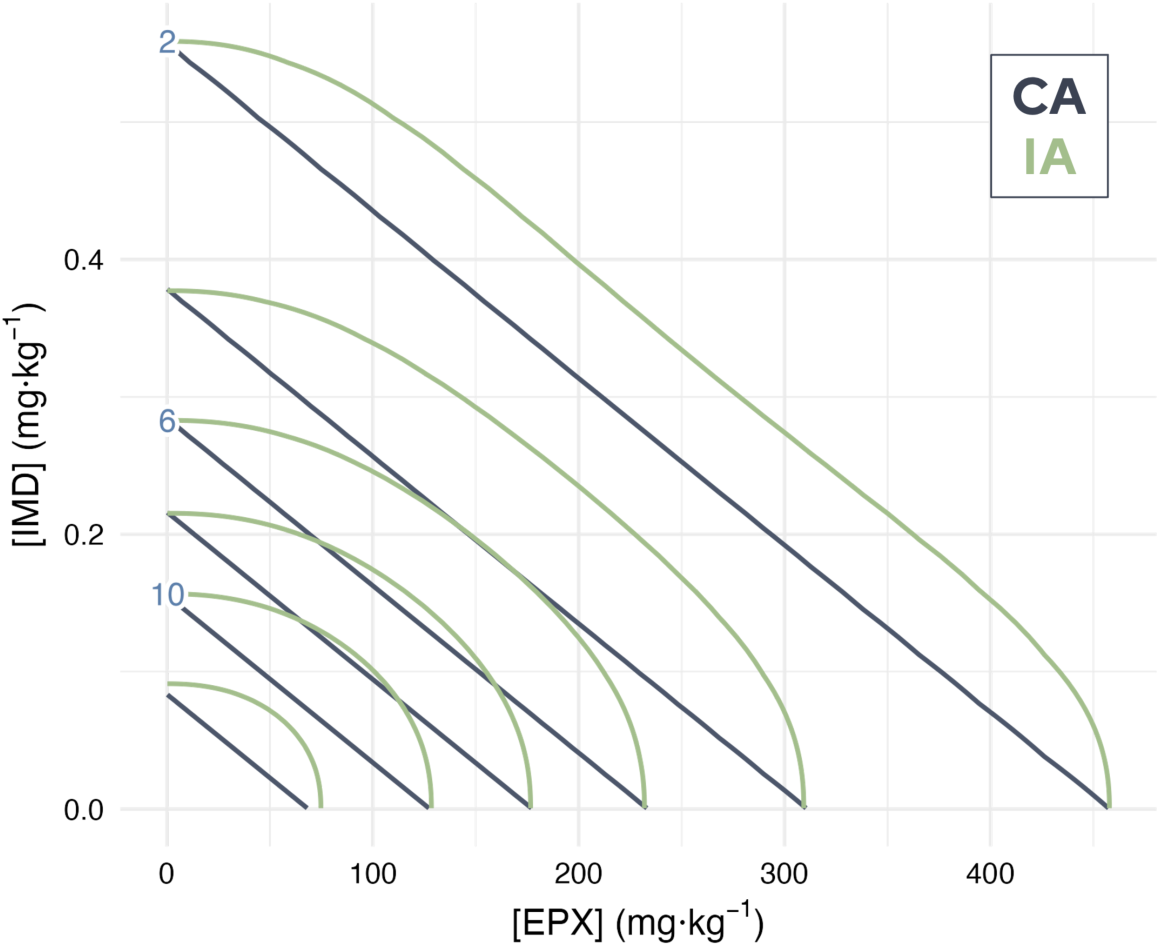
Estimated dose-response surfaced for cocoon production with the CA and IA model. Blue and green plain and dotted lines represent effect isoboles estimated with the CA model and the IA model, respectively. Numbers in blue indicate the predicted number of cocoons produced for a given isobole.

In contrast, the IA-SA model showed significant improvement over the independent action (IA) model (p = 0.0004) with an estimated interaction parameter a of −1.5 [−3.0 ; −0.2] (Figure 5). Consistently, the DL and DR models did not differ significantly from the SA model.

**Figure 5.**
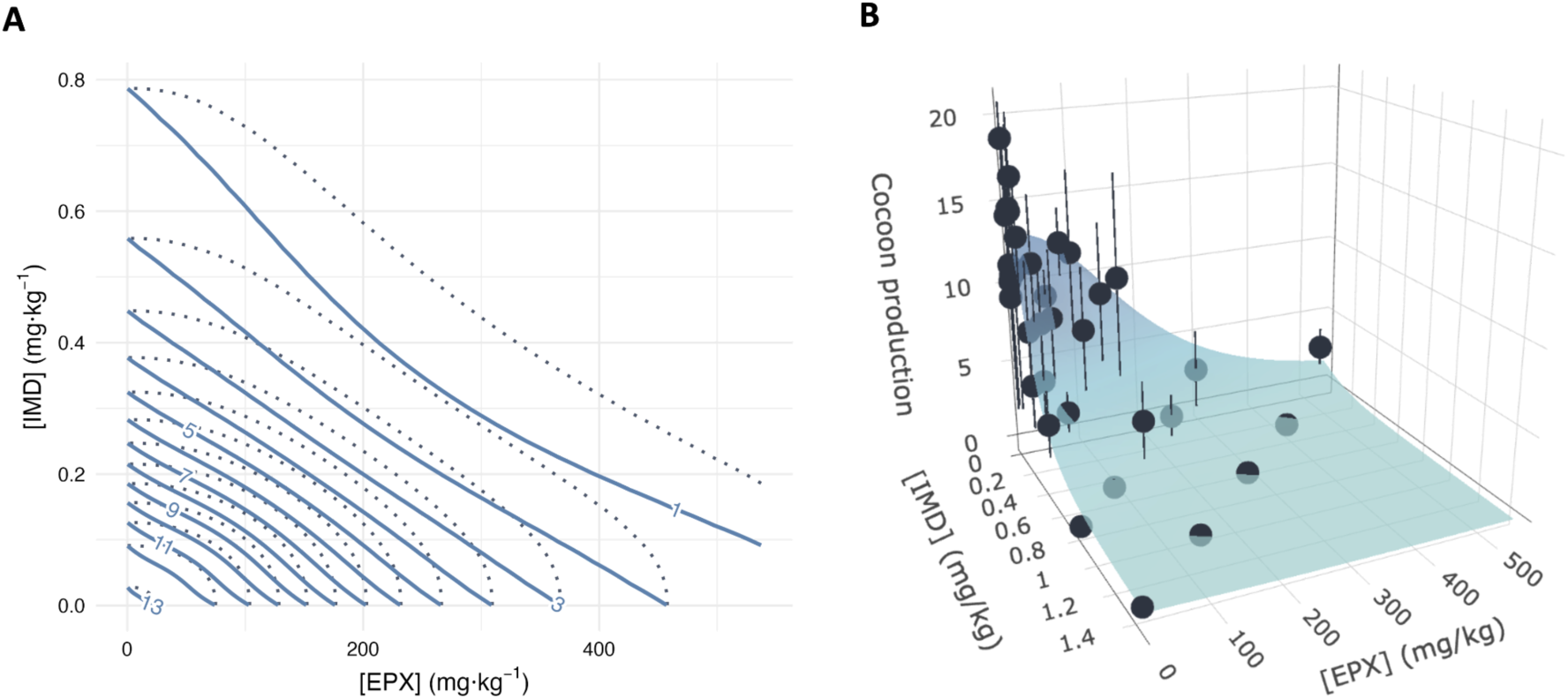
Estimated dose-response surface for cocoon production with the IA-SA model. A) Blue plain and dotted lines represent effect isoboles estimated with the IA-SA model and the IA model, respectively. B) The modeled dose-response surface with IA-SA is represented with means per condition (black bold dots) and their respective standard deviations (vertical black fine lines).

However, all lack-of-fit tests conducted for the CA and IA reference models, and their corresponding interaction models, were significant (Table S4.4). This indicates that the dose-response surfaces were not perfectly captured by any of the tested models.

## 4. Discussion

This study assessed the individual and combined effects of epoxiconazole and imidacloprid on the earthworm *Aporrectodea caliginosa*, with the expectation that imidacloprid would be more toxic and that cytochrome P450 inhibition by epoxiconazole could enhance its effects. The results confirmed the higher toxicity of imidacloprid, which strongly reduced both growth and reproduction. Analysis of the mixture response suggested a modest synergistic deviation from the independent action model under the tested conditions. Because of its conservative structure, the CA-SA model did not capture this deviation, whereas the IA-SA model did. However, significant lack-of-fit in the dose-response surface models indicates that these patterns should be interpreted with caution. Despite these limitations, the results suggest that IA-SA modeling provides a more accurate description of mixture effects under the tested exposure conditions and that synergistic interactions between insecticides and azole fungicides could represent an underestimated risk in soil ecotoxicology.

### 4.1. Single compound toxicity

The individual toxicity profiles of epoxiconazole and imidacloprid in earthworms confirm both compounds as environmentally relevant stressors, albeit with markedly different potencies. Imidacloprid demonstrated high toxicity to earthworms with pronounced effects on both growth (NOEC_juvenile_ = 0.28 mg/kg; NOEC_adult_ = 0.5 mg/kg) and reproduction (EC_50_ = 0.55 mg/kg; NOEC = 0.28 mg/kg). These findings align closely with published literature, where imidacloprid EC_50_ values for reproduction in *Eisenia* species have been reported at 0.87 mg/kg (Wang et al., 2019), and chronic assays showed EC_50_ values ranging from 0.21 to 1.89 mg/kg depending on soil type (Bandeira et al., 2020).

In contrast, epoxiconazole exhibited substantially lower toxicity, with moderate effects on growth (NOEC_juvenile_ = 9.3 mg/kg; NOEC_adult_ = 92.8 mg/kg) and reproduction (EC_50_ = 126.8 mg/kg; NOEC >= 200 mg/kg). The literature on epoxiconazole toxicity to earthworms remains more limited compared to imidacloprid, reflecting the generally lower concern for triazole fungicides in soil invertebrate risk assessment. The reduced potency measured in our study is consistent with reported data for *Aporrectodea icterica* exposed to a formulated version of epoxiconazole (OPUS®) in Pelosi et al. (2016), however the LC_50_ values of 45.5 mg/kg indicates a higher sensitivity of this species.

Across both compounds, a consistent pattern emerged: reproductive effects occurred at lower concentrations than adult growth effects, indicating that reproduction is a more sensitive endpoint for detecting pesticide impacts in adult earthworms. This sensitivity hierarchy aligns with established ecotoxicological principles and regulatory testing frameworks. However, juvenile growth was more sensitive than adult growth and at least as sensitive as reproduction, highlighting the particular vulnerability of early life stages.

The toxicity ratio between these compounds (EC_50_(IMD)/EC_50_(EPX) ≈ 230-fold difference) demonstrates the substantially higher potency of the neonicotinoid insecticide compared to the triazole fungicide, a pattern consistent with their respective modes of action and target specificity. This differential potency has important implications for mixture risk assessment, as the more toxic component (imidacloprid) would be expected to drive mixture effects under concentration addition assumptions.

### 4.2. Mixture effects - Observed results in accordance to theoretical expectations

A significant synergistic interaction in the epoxiconazole–imidacloprid mixture was detected relatively to the IA reference model (IA-SA model, *a* = −1.5), whereas the CA-SA model did not identify a departure from additivity, which is consistent with the more conservative nature of the CA reference. Power analysis confirmed that interactions of this magnitude (−1.5 or lower) were reliably detectable with the IA-SA model. This synergy suggested by the IA-SA model aligns with theoretical expectations based on toxicokinetic mechanisms described in the literature.

Importantly, retesting the single substances in the MIX experiment proved essential to avoid bias in the estimation of mixture effects. Although the dose-response curves from the UNI experiment were not statistically different, their estimated EC_50_ values differed substantially. Without retesting, this discrepancy could have generated an artificially strong synergistic signal (false positive), driven solely by the higher sensitivity of earthworms to imidacloprid in the second experiment rather than by a true mixture interaction.

However, a statistical limitation must be acknowledged: all lack-of-fit tests for the CA and IA reference models, as well as for their associated interaction models, showed significant lack-of-fit. Strictly speaking, this indicates that none of the fitted dose–response surfaces fully captured the variability in the data. While part of this lack-of-fit may be influenced by heterogeneous variance in the response, it cannot be fully attributed to the choice of error distribution, as alternative formulations (e.g., negative-binomial or quasi-Poisson) did not resolve the lack-of-fit. Significant deviations may therefore also reflect limitations in the mean structure of the fitted surfaces or biological processes not explicitly represented in the Jonker framework, such as nonlinear or context-dependent interactions. Consequently, interaction estimates derived from these models should be interpreted with caution. Additional replication or refined experimental designs would be required to more robustly characterize the magnitude and consistency of the observed synergistic pattern.

Nevertheless, the observed synergistic interaction is in accordance with the hypothesized toxicokinetic mechanism with distinct modes of action but with a metabolic inhibition of imidacloprid biotransformation by epoxiconazole through cytochrome P450 interference (Bass et al., 2015 ; Cedergreen et al., 2006 ; Puinean et al., 2010). Such metabolic interactions have been demonstrated in aquatic invertebrates such as *Daphnia magna*, where triazole co-exposure enhanced esfenvalerate toxicity up to fourfold (Cedergreen et al., 2006) and increased α-cypermethrin toxicity threefold (Gottardi et al., 2017). Similar synergistic interactions between atrazine and chlorpyrifos have also been reported in invertebrates, consistent with metabolic inhibition mechanisms. Additionally, the moderate hydrophilicity of imidacloprid (log K_ow_ ≈ 0.57) suggests that direct excretion pathways may predominate over metabolic transformation in earthworms, reducing the potential for metabolic interference which could explain the absence of strong interaction. Importantly, this metabolic mechanism remains unconfirmed in earthworms. Earthworms possess unique physiological adaptations for soil environments, including specialized detoxification mechanisms that may not rely heavily on cytochrome P450-mediated metabolism of imidacloprid. Earthworms possess detoxification strategies that rely on glutathione-S-transferases, carboxylesterases, antioxidant enzymes, and gut microbiota metabolism (Katagi & Ose, 2015; Wang et al., 2019). Evidence for imidacloprid metabolism via cytochrome P450 enzymes comes primarily from insect species (Chen et al., 2018; Chen et al., 2019), where cytochrome P450 overexpression has been linked to imidacloprid resistance (Elzaki et al., 2017; Kaplanoglu et al., 2017; Liang et al., 2015). However, the extent to which these metabolic pathways operate in earthworms remains largely unexplored, and extrapolation across taxonomically distant invertebrate groups may not be justified. Therefore, while the statistical signal points toward a small synergistic interaction, biological confirmation of the underlying mechanism is currently lacking.

These findings illustrate the difficulty of extrapolating both mechanisms and magnitudes of interaction across taxa, and emphasize the need for species-specific validation. Future work, including toxicokinetic measurements, or the integration of biochemical biomarkers such as catalase and glutathione-S-transferases as phase II detoxification enzymes (Givaudan et al., 2014; Pelosi et al., 2016), would be necessary to confirm whether the interaction detected here reflects a true synergistic process or arises from statistical noise due to overdispersion. Such complementary data would greatly strengthen mechanistic interpretation of mixture effects in earthworms.

### 4.3. Environmental relevance and risk assessment implications

Comparison of environmental concentrations with observed effect thresholds reveals important risk assessment insights. For imidacloprid, the reproduction EC_50_ of 0.55 mg/kg represents concentrations 36.4 to 189.6-fold higher than typical median environmental levels (0.0029-0.0151 mg/kg), but only 3.4-fold higher than maximum detected concentrations (0.16 mg/kg) (Table 3). The growth NOEC of 0.28 mg/kg similarly falls within 1.8-fold of maximum environmental concentrations. This suggests that while effects are unlikely under typical exposure scenarios, peak contamination events could approach concentrations of ecological concern. For epoxiconazole, the reproductive EC_50_ of 126.8 mg/kg provides larger safety margins, being 448-fold higher than maximum detected environmental concentrations (0.283 mg/kg) and over 3,000-fold higher than median concentrations (Table 3). The growth NOEC of 9.3 mg/kg maintains a 33-fold safety margin even against maximum environmental levels.

**Table 3.**
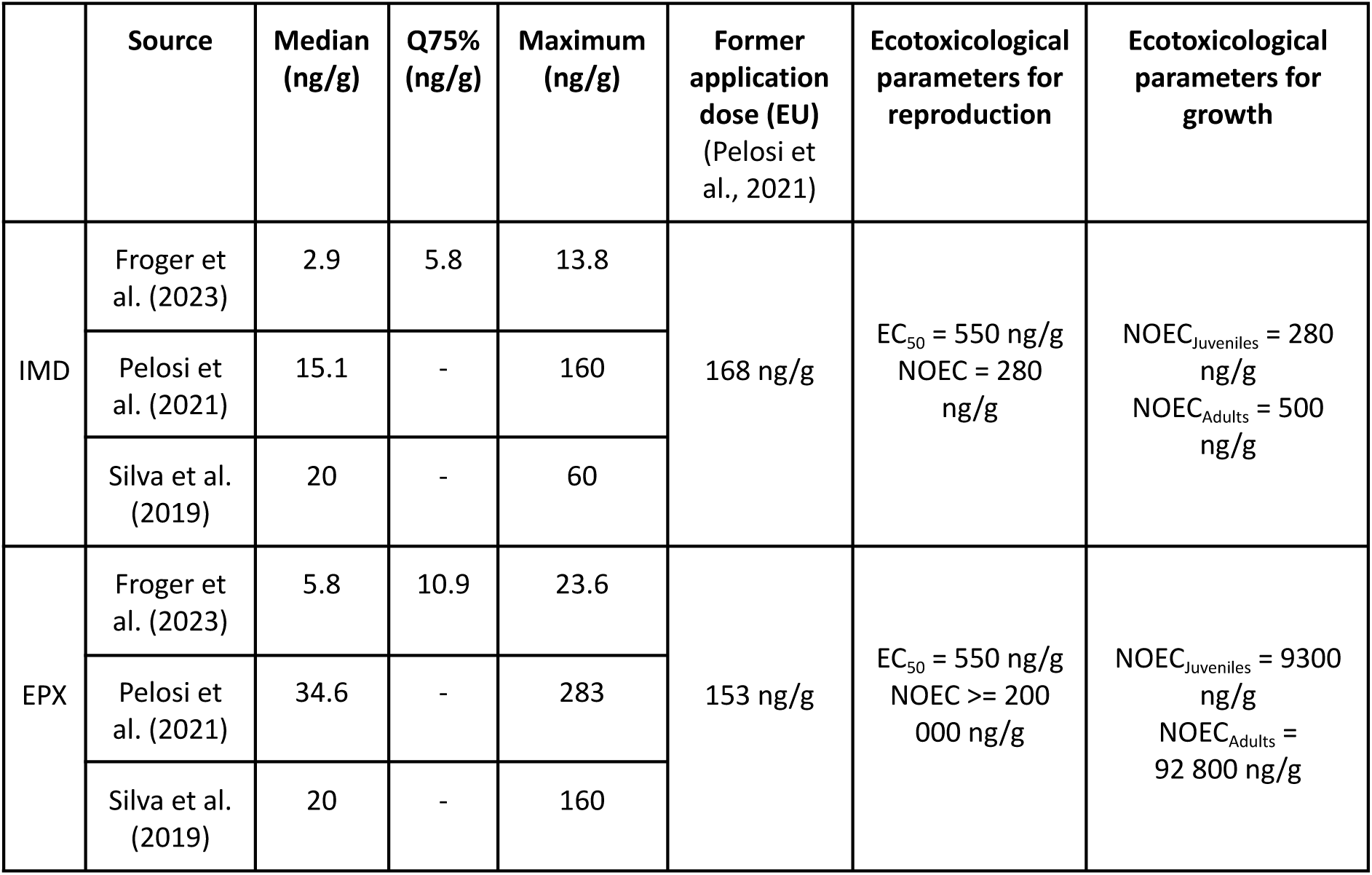
Summary of results for the detected imidacloprid and epoxiconazole residues in the Froger et al. (2023), Pelosi et al. (2021) and Silva et al. (2019) studies.

|  | Source | Median (ng/g) | Q75% (ng/g) | Maximum (ng/g) | Former application dose (EU) (Pelosi et al., 2021) | Ecotoxicological parameters for reproduction | Ecotoxicological parameters for growth |
| --- | --- | --- | --- | --- | --- | --- | --- |
| IMD | Froger et al. (2023) | 2.9 | 5.8 | 13.8 | 168 ng/g | EC<sub>50</sub> = 550 ng/g<br>NOEC = 280 ng/g | NOEC<sub>Juveniles</sub> = 280 ng/g<br>NOEC<sub>Adults</sub> = 500 ng/g |
|  | Pelosi et al. (2021) | 15.1 | - | 160 |  |  |  |
|  | Silva et al. (2019) | 20 | - | 60 |  |  |  |
| EPX | Froger et al. (2023) | 5.8 | 10.9 | 23.6 | 153 ng/g | EC<sub>50</sub> = 550 ng/g<br>NOEC ≥ 200 000 ng/g | NOEC<sub>Juveniles</sub> = 9300 ng/g<br>NOEC<sub>Adults</sub> = 92 800 ng/g |
|  | Pelosi et al. (2021) | 34.6 | - | 283 |  |  |  |
|  | Silva et al. (2019) | 20 | - | 160 |  |  |  |

These findings indicate that imidacloprid poses greater environmental risk than epoxiconazole under current contamination scenarios, particularly in intensive agricultural systems where maximum concentrations approach effect thresholds. The narrow safety margin for imidacloprid (1.8-3.4 fold) falls below typical regulatory safety factors of 10-100, suggesting potential for ecological impacts during peak exposure periods or in highly contaminated sites. Although both imidacloprid and epoxiconazole are now restricted, these substances are persistent in soils, with imidacloprid concentrations in some locations exceeding safety margins. Moreover, even though epoxiconazole alone exhibits limited toxicity, additive effects with imidacloprid residues could increase the overall risk to soil organisms.

Our results suggest that the combined effect of the two compounds could exceed what would be expected from the IA model. In this context, concentration addition (CA) remains a useful and precautionary reference, but it may still underestimate mixture toxicity when synergism occurs. The possibility of such synergistic deviation highlights that mixture exposures could pose greater risks than predicted from single-compound assessments alone. Consequently, our findings reinforce the importance of explicitly accounting for mixture effects and potential synergistic interactions in environmental risk assessment frameworks, especially for pesticide combinations that are likely to co-occur in soils.

### 4.4. Experimental Considerations and Resource Requirements

This mixture toxicity experiment involved 388 individual earthworms and required approximately 32 kg of soil. Such an experimental design illustrates the substantial logistical challenges commonly encountered in multi-stress studies on soil fauna. The need to handle large amounts of substrate and maintain individual organisms under controlled conditions necessitates considerable organizational, material, and human resources. These constraints also emphasize the importance of prioritizing target compounds for testing, based on their environmental relevance and the feasibility of experimental implementation.

The scale and complexity of such experiments highlight key strategic considerations for mixture toxicity research. Because they require substantial investments in time, personnel, materials, and laboratory capacity, comprehensive mixture studies cannot realistically be performed for every pesticide combination found in the environment. Instead, such resource-intensive approaches should be directed toward mixtures with strong mechanistic justification for interactions, clear environmental relevance, or specific regulatory significance. In this study, the EPX–IMD combination fulfilled these criteria due to its representative occurrence in current arable soil contamination profiles and documented potential for synergistic effects through cytochrome P450 metabolic interference.

This resource intensity highlights the need for strategic prioritization in mixture toxicity research, emphasizing the importance of preliminary screening approaches, computational modeling, and literature-based interaction predictions to identify the most promising candidates for comprehensive experimental investigation. Our study’s ability to detect a significant interaction provides important results that help refine our understanding of mixture effect predictions and guide future research prioritization. Such comprehensive studies, despite their demands, remain essential for validating theoretical interaction frameworks and ensuring robust environmental risk assessment approaches for pesticide mixtures in soil ecosystems.

## 5. Conclusion

This study provides a comprehensive evaluation of the mixture toxicity of epoxiconazole and imidacloprid in earthworms, revealing indications of synergistic interactions consistent with theoretical expectations based on cytochrome P450-mediated metabolic interference. Individual toxicity assessments confirmed both compounds as environmentally relevant stressors: imidacloprid exhibited high potency on growth (NOEC = 0.28 mg/kg) and reproduction (EC50 = 0.55 mg/kg), while epoxiconazole produced moderate effects (growth NOEC = 9.3 mg/kg; reproduction EC50 = 126.8 mg/kg). Reproductive endpoints were more sensitive than growth for both pesticides, in agreement with established ecotoxicological patterns.

Environmental risk assessment highlighted concerning exposure scenarios, particularly for imidacloprid, where maximum measured soil concentrations (0.16 mg/kg) approach effect thresholds, leaving safety margins of only 1.8-3.4 fold, values lower than typical regulatory assessment factors. The potential for synergistic interactions with epoxiconazole further increases the likelihood that combined exposures may surpass toxicologically relevant limits, underscoring the need to incorporate mixture considerations into risk evaluations. At the same time, the experimental resources required for robust mixture testing, especially when including factorial designs, emphasize the necessity of strategic prioritization in mixture toxicity research.

A remaining critical knowledge gap concerns whether the hypothesized metabolic interference mechanisms actually occur in earthworm systems. Given the significant lack-of-fit observed in our mixture models and the overdispersion in the intermediate values of our count data, the detected synergistic interaction should be interpreted cautiously. Additional data, such as increased replication, targeted toxicokinetic measurements, or independent mechanistic validation, are needed to confirm the biological basis and magnitude of the interaction.

To address this, dedicated toxicokinetic studies are required to quantify bioaccumulation dynamics and evaluate the plausibility of P450-related metabolic inhibition in earthworms. The integration of biochemical biomarkers (e.g., oxidative stress markers, glutathione-S-transferases) could provide complementary insight into underlying mechanisms. Ultimately, coupling such mechanistic information with Dynamic Energy Budget–ToxicoKinetic–ToxicoDynamic (DEB-TKTD) models will enable prediction of internal exposure and biological effects under realistic mixture scenarios, supporting more robust and process-based environmental risk assessment in agricultural soils.

## Author contribution statement (Credit syntaxe)

L. Gollot : Conceptualization, Data curation, Formal Analysis, Investigation, Methodology, Software, Validation, Visualization, Writing – Original Draft Preparation.

C. Tebby : Formal Analysis, Methodology, Resources, Software

L. Frattaroli : Data curation, Investigation, Methodology, Writing – Review & Editing.

R. Beaudouin : Formal Analysis, Funding Acquisition, Methodology, Resources, Supervision, Writing – Review & Editing.

R. Royauté : Conceptualization, Funding Acquisition, Methodology, Project Administration, Resources, Supervision, Writing – Review & Editing.

J. Faburé : Funding Acquisition, Methodology, Resources, Supervision, Writing – Review & Editing.

## Data statement

All data and R scripts used in this study are publicly available on GitHub (https://github.com/LisaGllt/Ew-Mix-TD) with an accompanying website providing comprehensive documentation and in-depth explanations (https://lisagllt.github.io/Ew-Mix-TD).

## Funding

This work was supported by the French National Research Agency (ANR) through the project EEWORM (N°ANR-23-CE34-0002); and the Biosphera graduate school of the University Paris-Saclay (PhD Grant).

## Disclaimer

The authors declare that they have no known competing financial interests or personal relationships that could have appeared to influence the work reported in the present study.

## Acknowledgments

The authors thank Cheikh Ahmed Tidiane Sarr, Julien Rouyer, Antoine Bamière and Véronique Etievant for their technical support.

## Notes

https://github.com/LisaGllt/Ew-Mix-TD

### Competing Interest Statement

The authors have declared no competing interest.

