## Supplemental Materials for "Toxic cocktails in soils - Evidence for synergistic effects of the imidacloprid-epoxiconazole mixture on earthworm life-history traits"

### Toxic cocktails in soils : Assessing fungicide-insecticide mixture effects on earthworm growth and reproduction

**References** **35**

4.1 Dose-response curves obtained for each mixture ratio. The lines represent the dose-response curves obtained (with common $Y_{max}$ and $Y_{min}$ set to 0). Crosses correspond to the number of cocoons produced by one cosm and the bold points represent the mean per condition. . 27
4.2 Estimated dose-response surfaces isoboles by the CA (grey) and the IA (blue) model. Blue numbers aside each isobole correspond to the level of the isobole in produced cocoons. . . 30

4.3 Estimated dose-response surfaces isoboles by the IA-SA (blue) model. The isoboles from the IA model are represented in dotted lines. . . . . . . . . . . . . . . . . . . . . . . . . 31
4.4 Results of the CA model fit on experimental data. Thin black and grey lines represent the 2-folds and 5-fold changes, respectively. The thicker black line represents the identity line. 32
4.5 Results of the IA model fit on experimental data. Thin black and grey lines represent the 2-folds and 5-fold changes, respectively. The thicker black line represents the identity line. 32
4.6 Results of the CA-SA model fit on experimental data. Thin black and grey lines represent the 2-folds and 5-fold changes, respectively. The thicker black line represents the identity line. . . . . . . . . . . . . . . . . . . . . . . . . . . . . . . . . . . . . . . . . . 33
4.7 Results of the IA-SA model fit on experimental data. Thin black and grey lines represent the 2-folds and 5-fold changes, respectively. The thicker black line represents the identity line. . . . . . . . . . . . . . . . . . . . . . . . . . . . . . . . . . . . . . . . . . 33

### List of Tables

1.1 Main characteristics of the breeding soil from Versailles. Table from Bart et al. (2017). . . 6

2.1 NOEC and LOEC estimated with a Dunnett test. . . . . . . . . . . . . . . . . . . . . . . . 12
2.2 Estimated parameters of the dose-response curve for cocoon production of earthworms exposed to imidacloprid or epoxiconazole and their respective 95% confidence intervals. . 13
2.3 ANOVA table . . . . . . . . . . . . . . . . . . . . . . . . . . . . . . . . . . . . . 13
2.4 NOEC and LOEC estimated with a Dunnett test. . . . . . . . . . . . . . . . . . . . . . . . 13
2.5 NOEC and LOEC estimated with a Dunnett test. . . . . . . . . . . . . . . . . . . . . . . . 14
2.6 Estimated parameters of the dose-response curve for normalized cocoon production of earthworms exposed to imidacloprid or epoxiconazole and their respective 95% confidence intervals (with common slope and Ymax). . . . . . . . . . . . . . . . . . . . . . . . . 14
2.7 Estimated parameters of the dose-response curves from experiment B (common slope and $Y_{max}$, used in Jonker models. . . . . . . . . . . . . . . . . . . . . . . . . . . . . . 16

3.1 Power, misclassification and estimation accuracy of the SA model across interaction strengths under the CA and IA reference models. . . . . . . . . . . . . . . . . . . . . . . 26

4.1 Estimated dose-response curves for each mixture ratios. $Y_{min}$ set to 0. . . . . . . . . . . . 28
4.2 Dose-response curves estimated with a different slopes. . . . . . . . . . . . . . . . . . . 28
4.3 ANOVA-like table . . . . . . . . . . . . . . . . . . . . . . . . . . . . . . . . . . . 28
4.4 Estimated interaction parameters by the different Jonker interaction models. . . . . . . . 29
4.5 Comparison of the different models with Likelihood Ratio tests. . . . . . . . . . . . . . . . 29

### 1 Earthworm culture

#### 1.1 Breeding and experiment soil

Table 1.1: Main characteristics of the breeding soil from Versailles. Table from Bart et al. (2017).

| Characteristics | Value |
|---|---|
| Clay ( $<$ 2 µm, g/kg) | 226.0 |
| Fine silt (2-20 µm, g/kg) | 174.1 |
| Coarse silt (20-50 µm, g/kg) | 298.9 |
| Fine sand (50-200 µm, g/kg) | 239.1 |
| Coarse sand (200-2000 µm, g/kg) | 47.9 |
| $CaCO_3$ total (g/kg) | 23.3 |
| Organic matter (g/kg) | 32.6 |
| $P_2O_5$ (g/kg) | 0.1 |
| Organic carbon (g/kg) | 18.9 |
| Total nitrogen (N) (g/kg) | 1.5 |
| C/N | 12.5 |
| pH | 7.5 |
| Total $Cu$ (mg/kg) | 25.2 |

#### 1.2 DNA barcoding

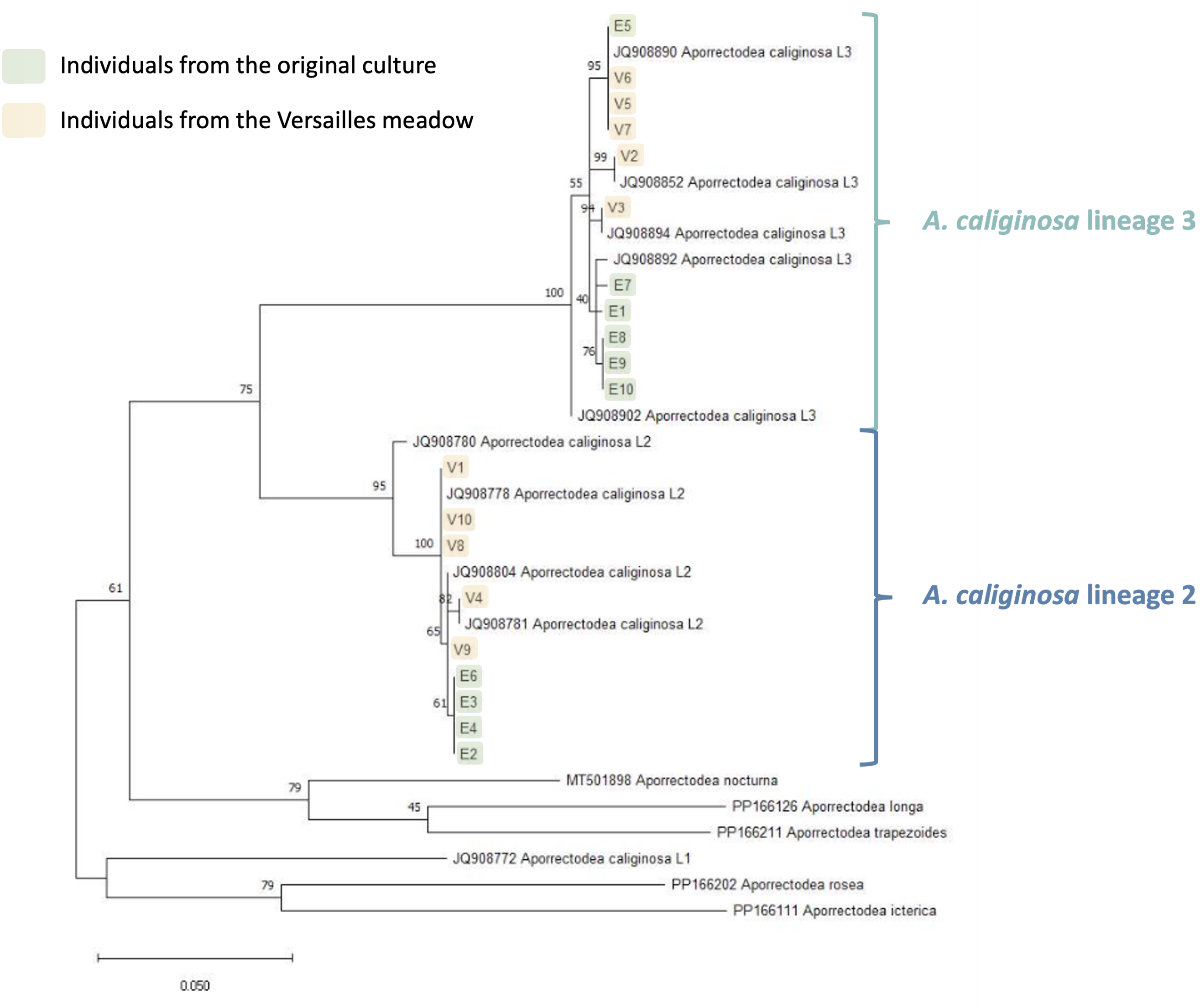

Figure 1.1: Results of the DNA barcoding of 10 individuals from the culture (highlighted in yellow)

### 2 Experiments

#### 2.1 Growth and reproduction experiments

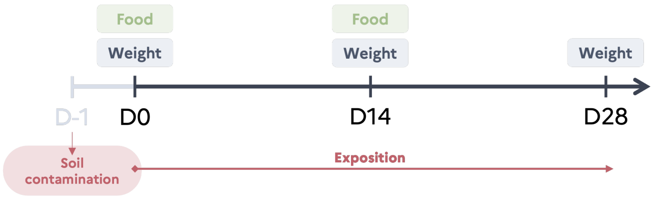

Figure 2.1: Overview of the growth experiments

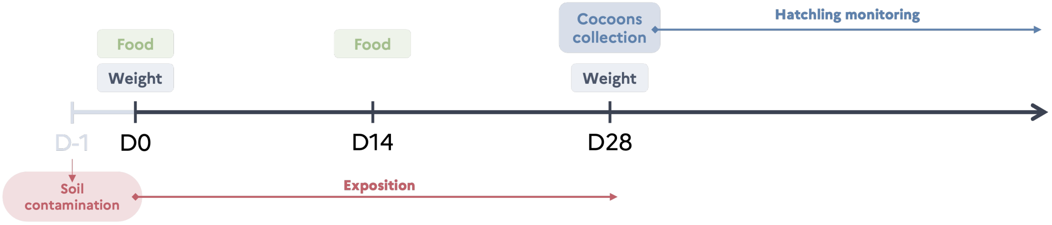

Figure 2.2: Overview of the reproduction experiments

#### 2.2 Experiment A - Single substance

##### 2.2.1 Juvenile growth

*Initial state of earthworms*

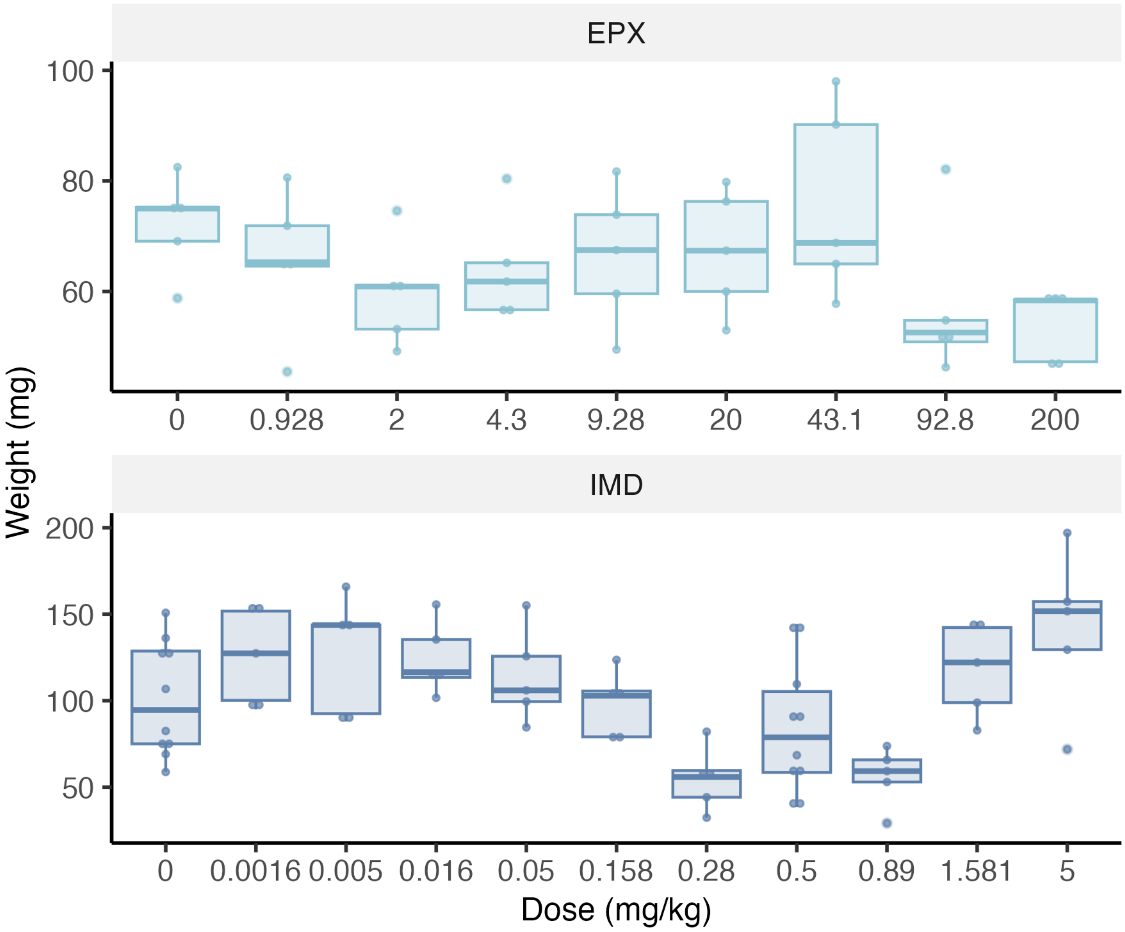

Figure 2.3: Initial weight of juvenile earthworms.

For IMD, there is a significant effect of the dose on earthworm initial weights.

- ANOVA : pval(IMD) = 9.66e-05,
- All concentrations are not significatively different from the control (post hoc Dunett test).

However, earthworms were non the less randomly attributed to conditions and the growth model allows to take the differences in initial weight into account.

For EPX, there is no significant effect of the dose on the initial weight of earthworms (Student test : pval(IMD) = 0.125).

***Mortality during the experiment***

Mortality was observed for the highest concentrations of IMD (Figure 2.4).

***Growth curves***

Growth curves of juveniles exposed to IMD or EPX are presented Figure 2.5a and Figure 2.5b

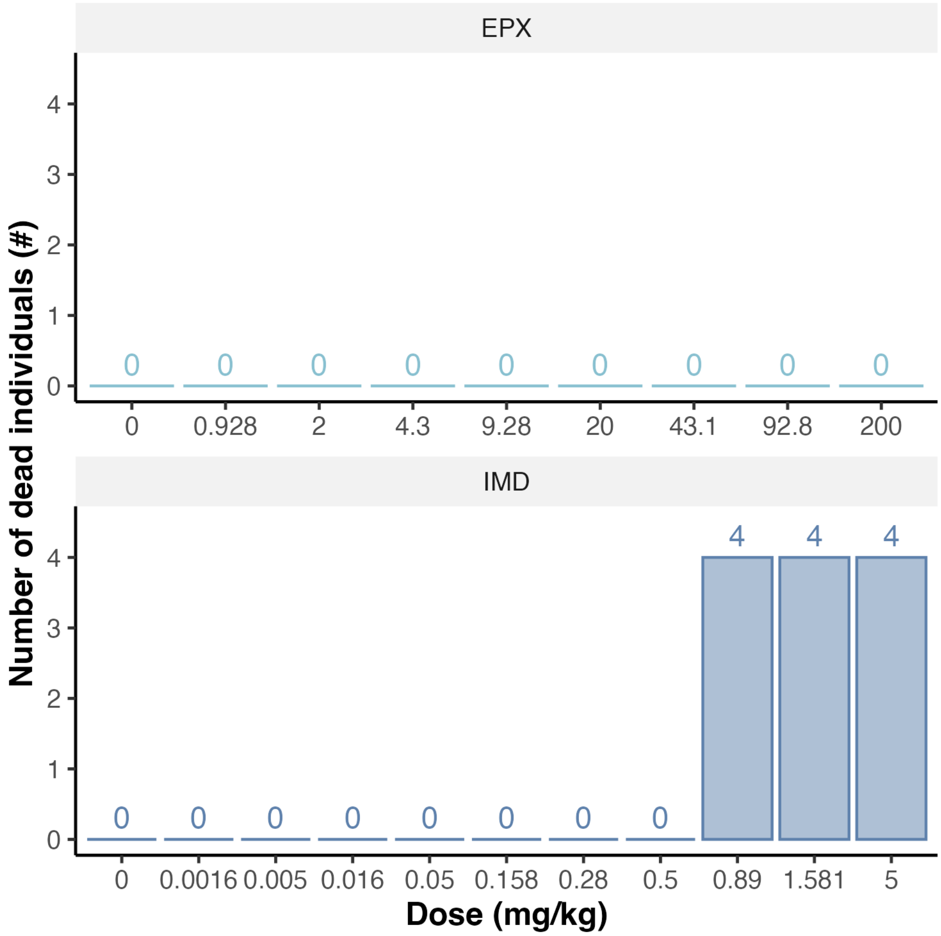

Figure 2.4: Number of dead individuals per condition at the end of the experiment

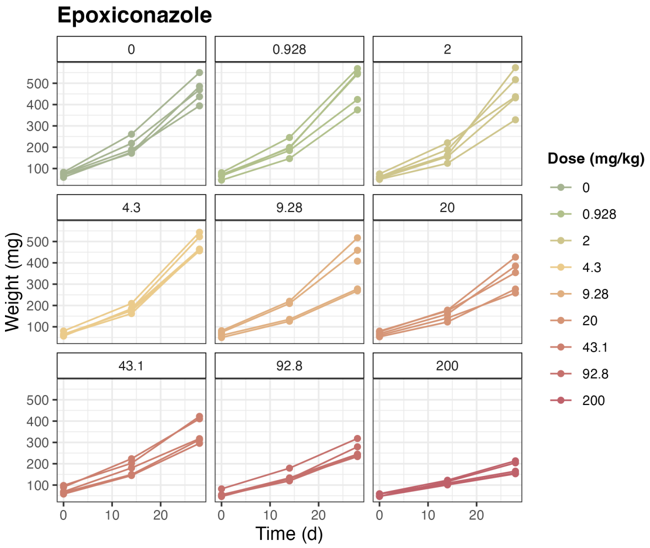

(a) Growth curves of juveniles exposed to EPX

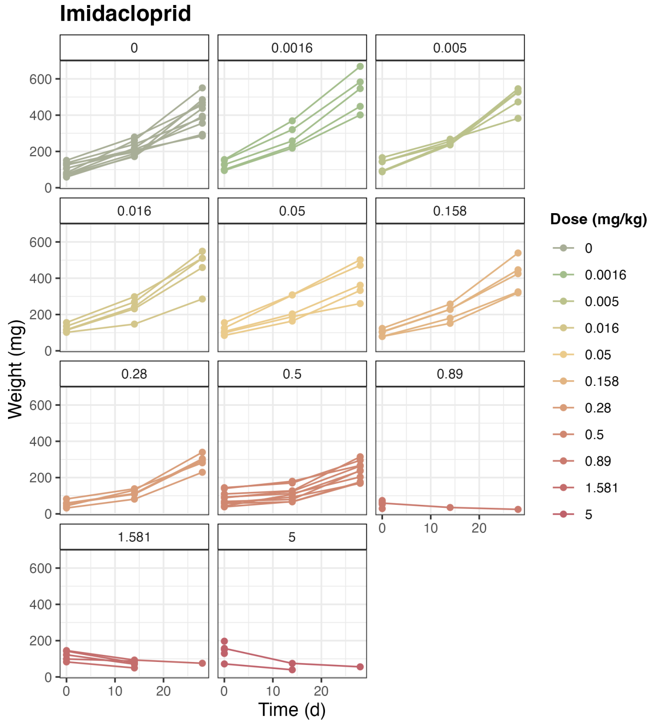

(b) Growth curves of juveniles exposed to IMD

Figure 2.5: Growth curves of juvenile earthworms.

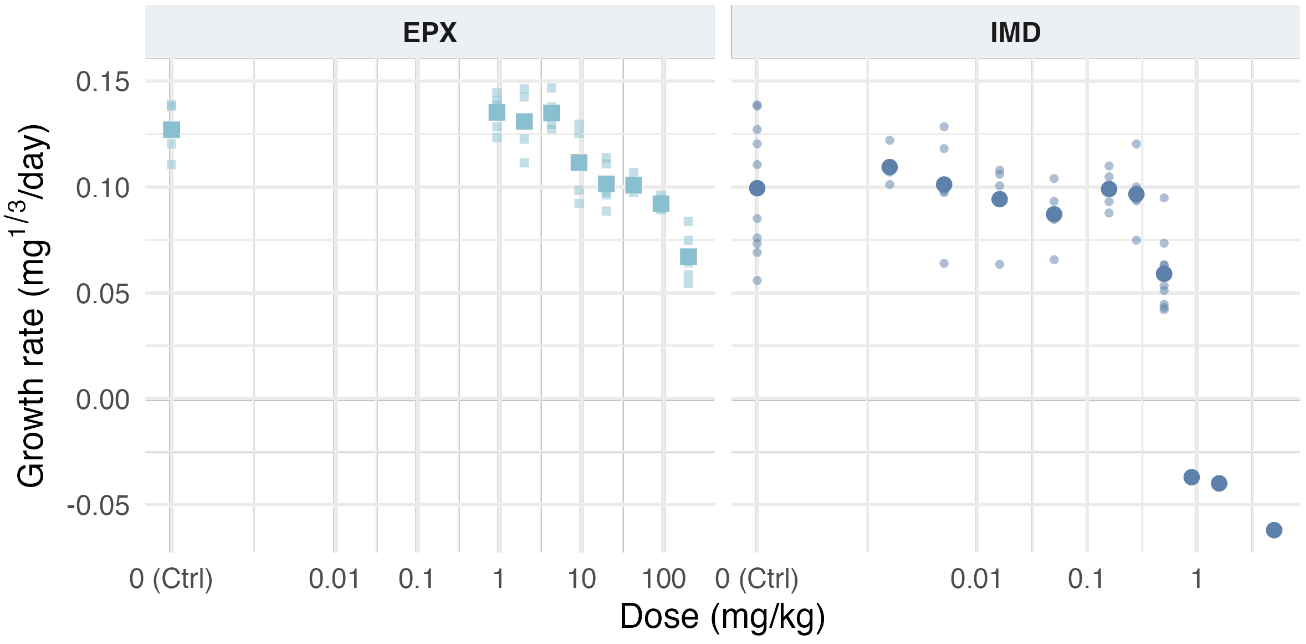

Figure 2.6: Effect of the pesticides on juvenile growth rates

Table 2.1: NOEC and LOEC estimated with a Dunnett test.

| | NOEC (mg/kg) | LOEC (mg/kg) |
|---|---|---|
| IMD | 0.28 | 0.5 |
| EPX | 9.28 | 20 |

##### 2.2.2 Reproduction

***Initial state***

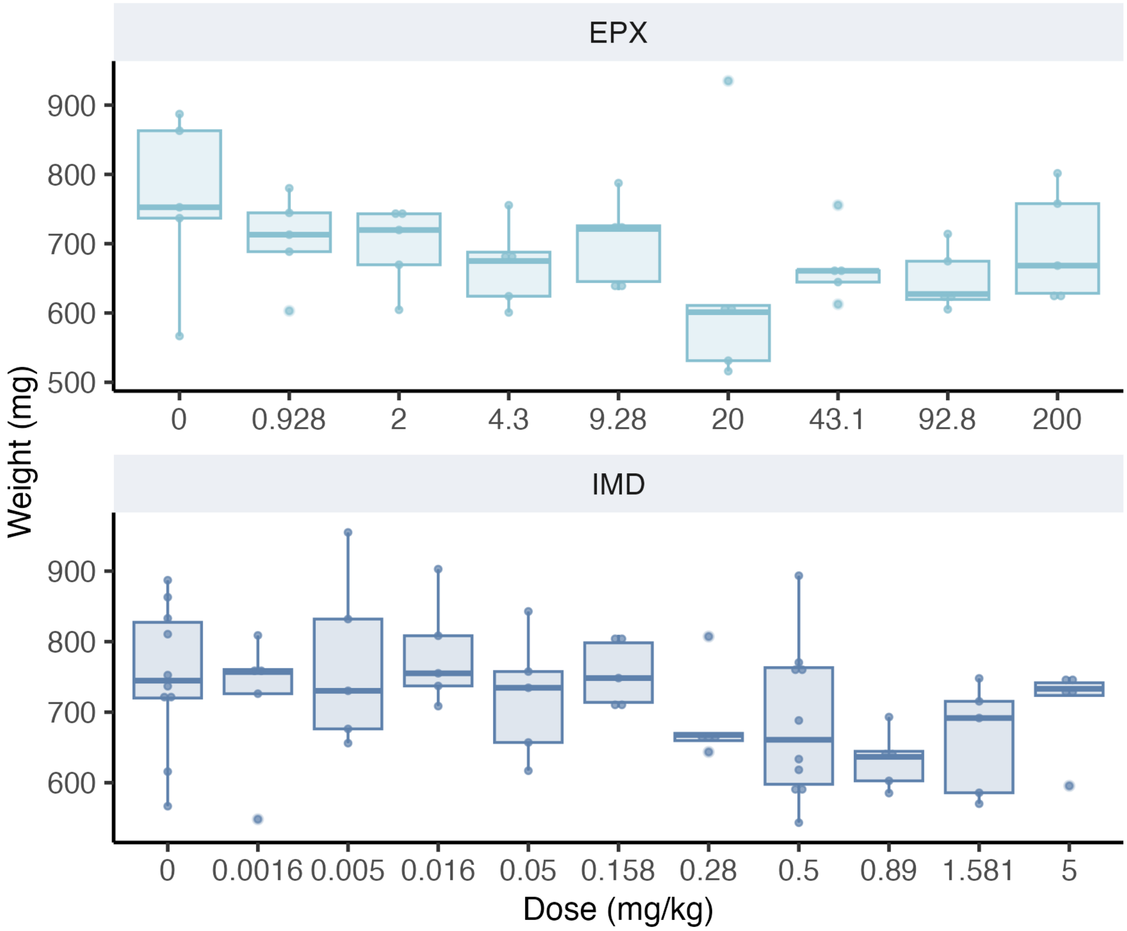

Figure 2.7: Initial weight of adult earthworms.

There is no effect of the dose on the initial weight of earthworms (Student test : pval(EPX) = 0.17 & pval(IMD) = 0.573).

***Effect on cocoon production***

Table 2.2: Estimated parameters of the dose-response curve for cocoon production of earthworms exposed to imidacloprid or epoxiconazole and their respective 95% confidence intervals.

| | Imidacloprid | Epoxiconazole |
|---|---|---|
| $Y_{min}$ | 0 (fixed) | 0 (fixed) |
| $Y_{max}$ [CI 95%] | 14.3 [13.0 ; 15.6] | 18.7 [16.8 ; 20.7] |
| $EC_{50}$ (mg/kg) | 0.55 [0.45 ; 0.65] | 126.8 [73.6 ; 180.0] |
| | 2.8 [1.8 ; 3.8] | 1.1 [0.5 ; 1.6] |

lacke-of-fit test : p value = 1.

Log-Likelihood ratio test - Comparison of the dose-response curve models with or without common slope with data normalized with respective controls :

Table 2.3: ANOVA table

| Model | Model Df | LogLik | Df | p value |
|---|---|---|---|---|
| Different slopes | 6 | -518.05 | 0 | |
| Common slope | 5 | -518.30 | 1 | 0.4803 |

Table 2.4: NOEC and LOEC estimated with a Dunnett test.

| | NOEC (mg/kg) | LOEC (mg/kg) |
|---|---|---|
| IMD | 0.28 | 0.5 |
| EPX | >=200 | >=200 |

***Effect on adult growth***

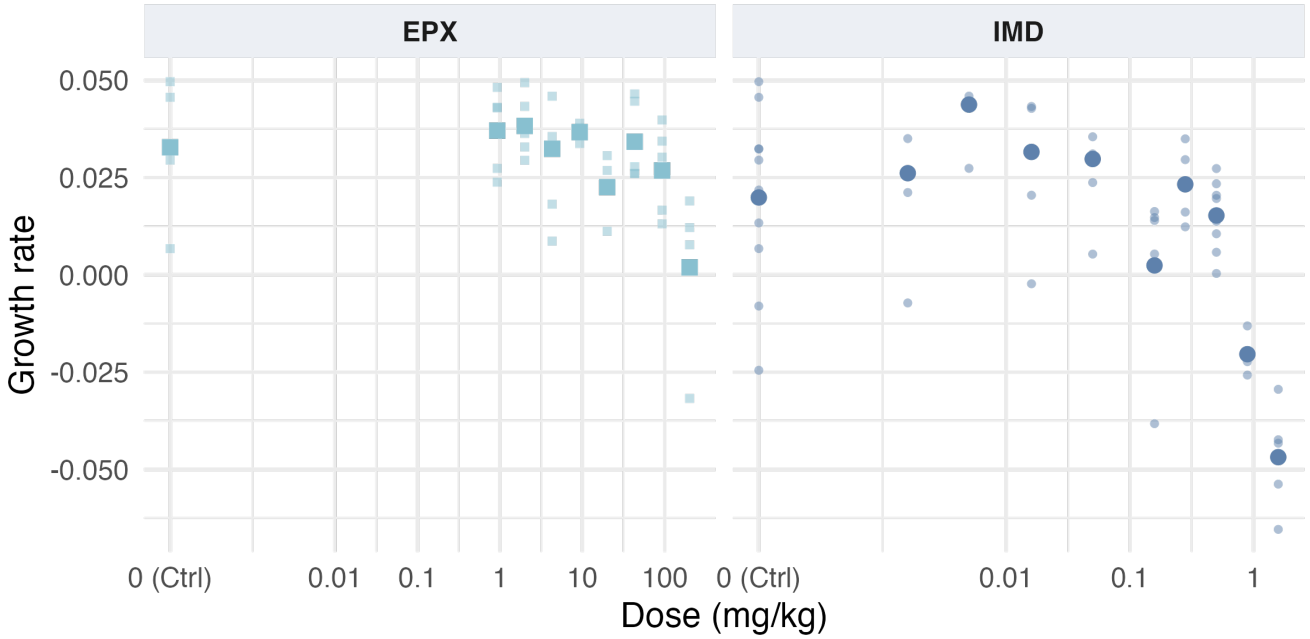

Figure 2.8: Effect of the pesticides on adult growth rates.

Table 2.5: NOEC and LOEC estimated with a Dunnett test.

| | NOEC (mg/kg) | LOEC (mg/kg) |
|---|---|---|
| IMD | 0.5 | 0.89 |
| EPX | 92.8 | 200 |

***Design determination***

Table 2.6: Estimated parameters of the dose-response curve for normalized cocoon production of earthworms exposed to imidacloprid or epoxiconazole and their respective 95% confidence intervals (with common slope and Ymax).

| | Imidacloprid | Epoxiconazole |
|---|---|---|
| $Y_{min}$ | 0 (fixed) | 0 (fixed) |
| $Y_{max}$ [CI 95%] | 107.9 [98.2 ; 117.6] | 107.9 [98.2 ; 117.6] |
| $EC_{50}$ (mg/kg) | 0.45 [0.27 ; 0.62] | 137 [57 ; 217] |
| | 1.55 [0.7 ; 2.4] | 1.55 [0.7 ; 2.4] |

#### 2.3 Experiment B - Mixture

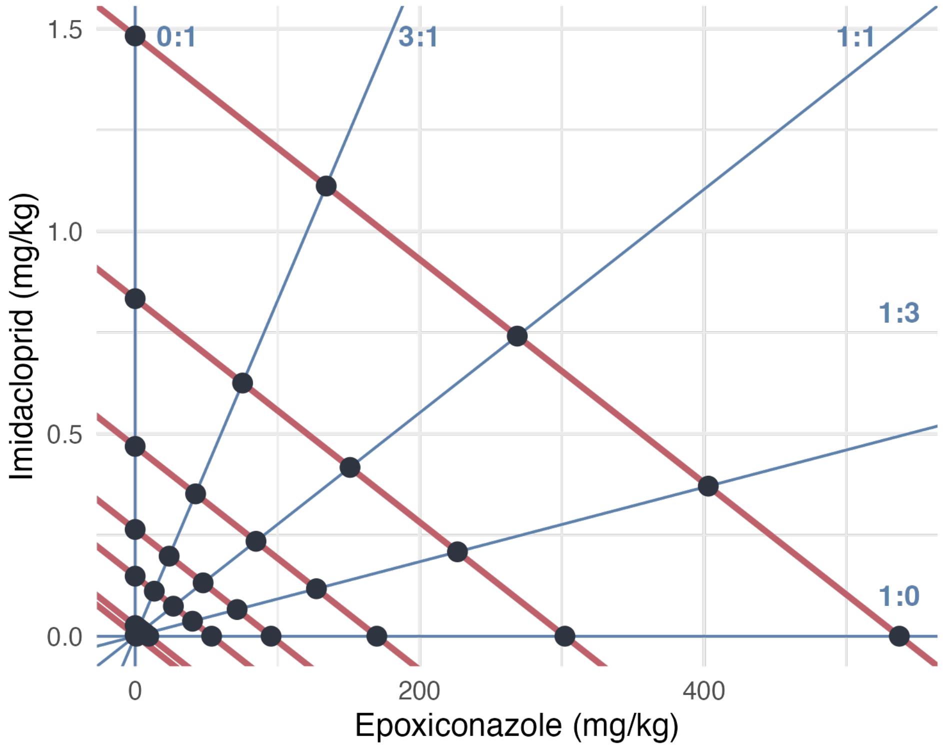

Figure 2.9: Experimental design of experiment B.

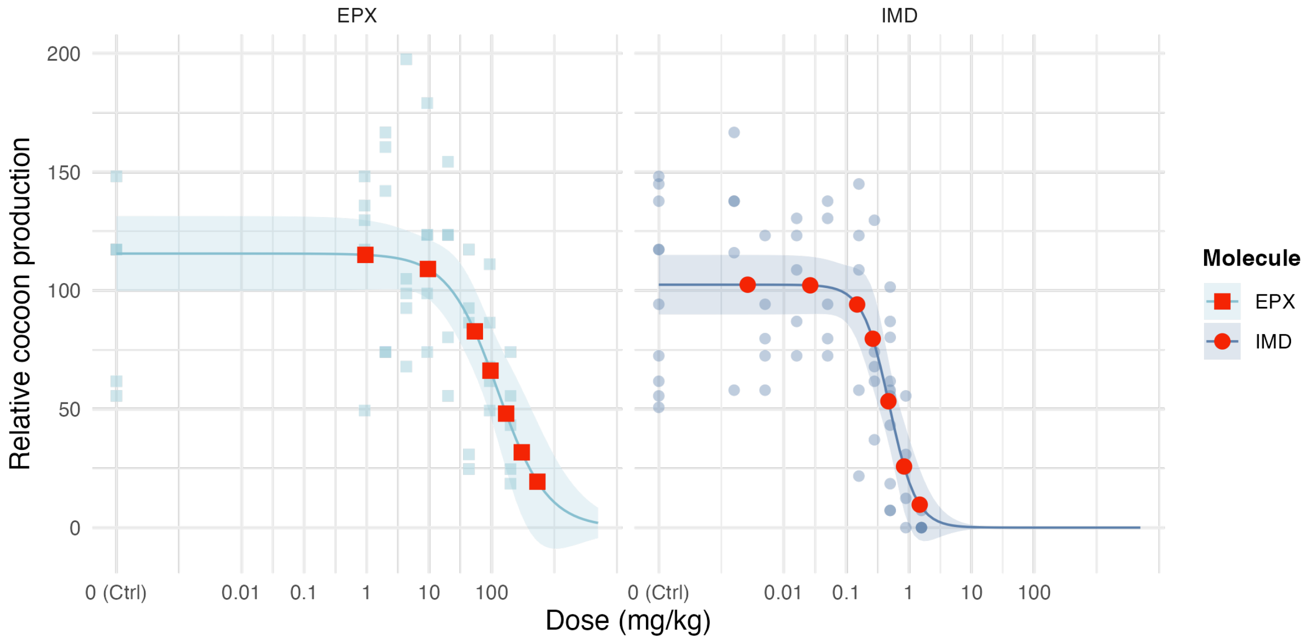

Figure 2.10: Expected effect range for each effect isobole (red points).

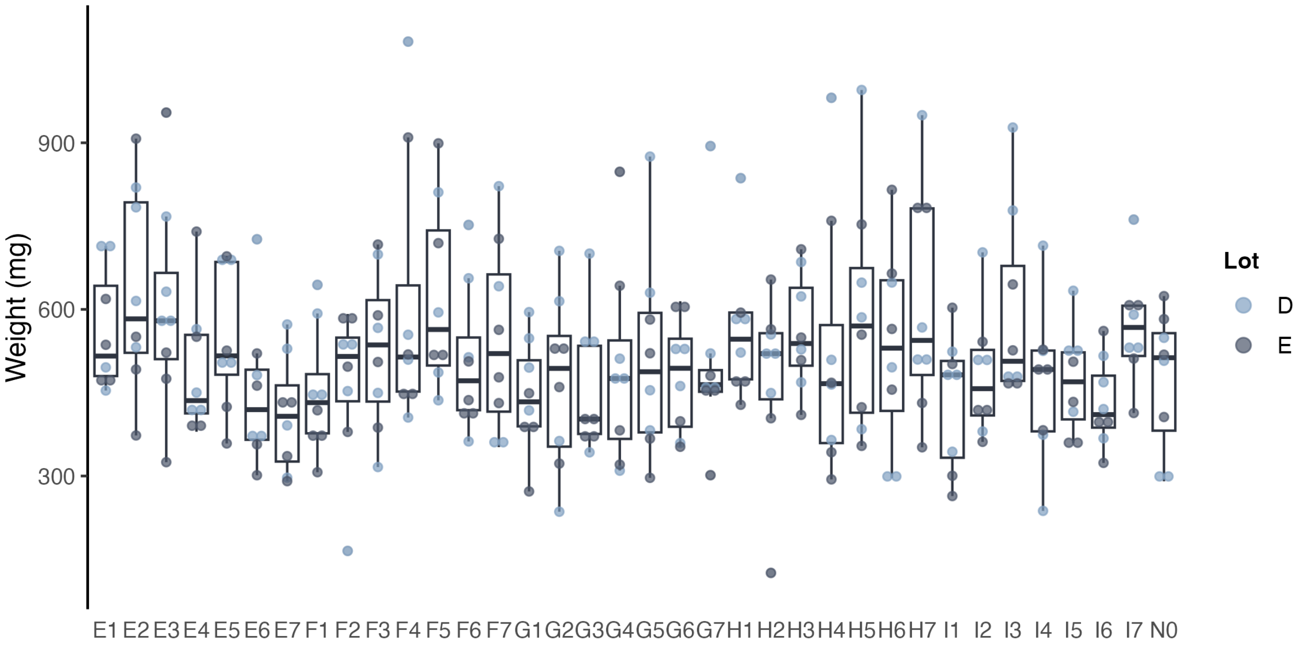

Figure 2.11: Initial body weight of earthworms used in the experiment B. Earthworm assigned to lot D are represented in blue while the ones assigned to lot E are represented in dark grey.

Table 2.7: Estimated parameters of the dose-response curves from experiment B (common slope and $Y_{max}$, used in Jonker models.

| | Imidacloprid | Epoxiconazole |
|---|---|---|
| $Y_{min}$ | 0 (fixed) | 0 (fixed) |
| $Y_{max}$ [CI 95%] | 13.1 [11.6 ; 14.5] | 13.1 [11.6 ; 14.5] |
| $EC_{50}$ (mg/kg) | 0.26 [0.20 ; 0.33] | 216 [160 ; 271] |
| | 2.28 [1.65 ; 2.9] | 2.28 [1.65 ; 2.9] |

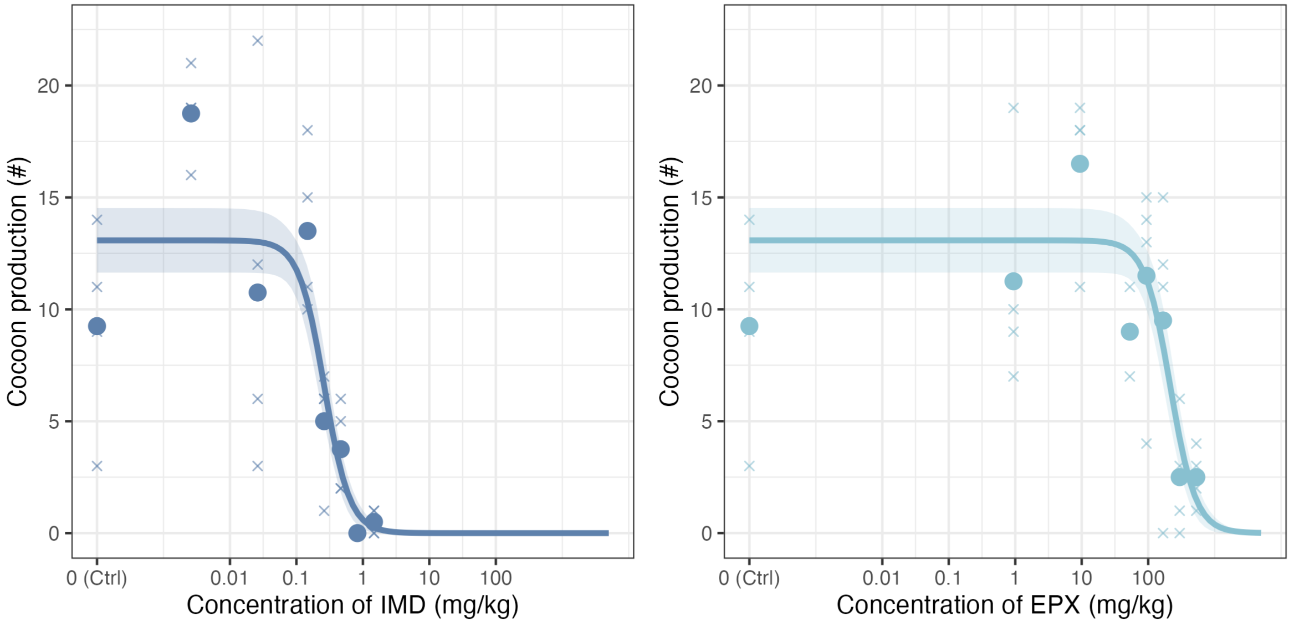

Figure 2.12: Dose response curves obtained with cocoon production. The line represents the modelled dose-response curve (with common $Y_{max}$ and slope for the 2 active substances). Crosses correspond to the number of cocoons produced by one cosm and the bold points represents the mean per dose.

### 3 Jonker interaction models

#### 3.1 Mathematical formulation

Jonker et al. (2005) describes a workflow to analyse mixture data.

##### 3.1.1 With CA as reference

Let $f_A$ and $f_B$ be the functions describing the dose-response curves of substance A and substance B, respectively.

In the case of no interaction, concentration addition implies the dose-response surface of the mixture of A and B can be expressed as all couple of concentration of substance A and substance B ($c_A$ ; $c_B$) as :

$$\frac{c_A}{f_A^{-1}(Y)} + \frac{c_B}{f_B^{-1}(Y)} = 1 \tag{3.1}$$

The Jonker simple interaction model add the definition of an interaction parameter ($a$) so the dose-response surface of the mixture of A and B can be expressed as all ($c_A$ ; $c_B$) as :

$$\frac{c_A}{f_A^{-1}(Y)} + \frac{c_B}{f_B^{-1}(Y)} = \exp(G) \tag{3.2}$$

With :

$$G(z_A, z_B) = a \times (z_A \times z_B) \tag{3.3}$$

And :

$$z_i = \frac{\mathrm{TU}_{x_i}}{\mathrm{TU}_{x_A} + \mathrm{TU}_{x_B}}, \quad \text{where} \quad \mathrm{TU}_{x_i} = \frac{c_i}{EC_{x_i}} \tag{3.4}$$

With :

- $Y$ : The response for a given couple of concentrations,
- $x$ : Usually 5, 10, ou 50 (50 used in this study),
- $z_A$ & $z_B$ : Contribution of each substance to the overall effect,

For the SA model, the interaction parameter value describes the type of the interaction :

- $a = 0$ : Additivity,
- $a < 0$ : Synergy,
- $a > 0$ : Antagonisme.

The Jonker framework adds two other types of model, making it four models in total :

- CA : Concentration addition, case with no interaction (Equation 3.1)
- SA : Simple interaction with constant deviation from additivity (Equation 3.3),
- DL : Dose-level dependent interaction where effects vary with response magnitude (Equation 3.5),
- DR : Ratio-dependent interaction where effects vary with mixture composition (Equation 3.5).

The DL and DR models include a second interaction parameter, $b$. The expression of $G$ is then :

$$G(z_A z_B) = \begin{cases} a\,[1 - b_{DL}]\, z_A z_B & \text{(DL model)} \\ (a + b_{DR} z_A)\, z_A z_B & \text{(DR model)} \end{cases} \tag{3.5}$$

##### 3.1.2 With IA as reference

Let $q_i$ be the function as $f_i(c_i) = Y_{max}\ q_i$.

With an endpoint inversely proportional to the dose, the Jonker interaction models can now be expressed as the following equations :

$$Y = Y_{max} \Phi(\Phi^{-1}[q_A(c_A) * q_B(c_B)] + G)$$

With :

- $Y$ : The response for a given couple of concentrations ($c_A$ and $c_B$),
- $Y_{max}$ : The maximum response (control response),
- $q_i(c_i)$ : Non-response probability of substance $i$,
- $\Phi$ : The standard cumulative normal distribution function.

And with :

$$G(z_A, z_B) = \begin{cases} a \times z_A z_B & \text{for the SA model} \\ a[1 - b \times q_A(c_A)\ q_B(c_B)]\ z_A z_B & \text{for the DL model} \\ (a + b\ z_A)\ z_A z_B & \text{for the DR model} \end{cases}$$

#### 3.2 Computational implementation

##### 3.2.1 With CA as reference

In practice, the fit of the dose-response curve surface is a double optimization :

1. For given set of interaction parameters, the corresponding effects must be found also numerically.
2. Numerical optimization of the interaction parameters

To perform step 1, a function `CA_complete2()` was written. For a given set of interaction parameters, finding the corresponding efect for a couple of concentration ($c_A$ ; $c_B$) implies the minimization of the following term :

$$\frac{c_A}{f_A^{-1}(Y)} + \frac{c_B}{f_B^{-1}(Y)} - \exp(G) \tag{3.6}$$

For step 2, a function `CA_complete_fit_speed()` was written. Parameter estimation for the interaction models requires maximum likelihood estimation (MLE). The optimization procedure depends on the assumed error distribution: for normally distributed errors, this is equivalent to minimizing the residual sum of squares (calculated with the function `CA_complete2_RSS()`), while for Poisson-distributed errors, the log-likelihood function (Equation 3.7) (calculated with the function `CA_complete2_Poisson()`) is directly maximized.

Let $\mathbf{y} = (y_1, y_2, \ldots, y_n)$ be the vector of observed responses,   the parameter vector to be estimated, and $f(.| )$ the density function conditional to the parameters. Then, the likelihood function is defined as :

$$\mathcal{L}(\theta|\mathbf{y}) = \prod_{i=1}^{n} f(y_i|\theta)$$

The log-likelihood function, which is computationally more tractable, is given by:

$$l(\theta|\mathbf{y}) = \ln(\mathcal{L}(\theta|\mathbf{y})) = \sum_{i=1}^{n} \ln(f(y_i|\theta))$$

For count data (cocoon production), assuming $y_i \sim \mathcal{P}(\lambda_i)$ where $\lambda_i = g(\mathbf{x_i}; \theta)$ represents the expected response given covariates $\mathbf{x_i}$, the log-likelihood becomes:

$$\begin{aligned} l(\theta|\mathbf{y}) &= \sum_{i=1}^{n} \ln\left(\frac{e^{-\lambda_i}\lambda_i^{y_i}}{y_i!}\right) \\ &= \sum_{i=1}^{n} [y_i \ln(\lambda_i) - \lambda_i - \ln(y_i!)] \\ &= \sum_{i=1}^{n} [y_i \ln(g(x_i;\theta)) - g(x_i;\theta) - \ln(y_i!)] \end{aligned} \tag{3.7}$$

##### 3.2.2 With IA as reference

It follows the same principle but the calculation of the effect of the mixture is more straight forward.

#### 3.3 Power analysis

**Probability of direction (pd)** The probability of direction is the posterior probability that a model parameter has a positive or a negative sign. It is defined as the proportion of the posterior distribution that lies on the same side of zero as the median, and quantifies the certainty about the direction of an effect.

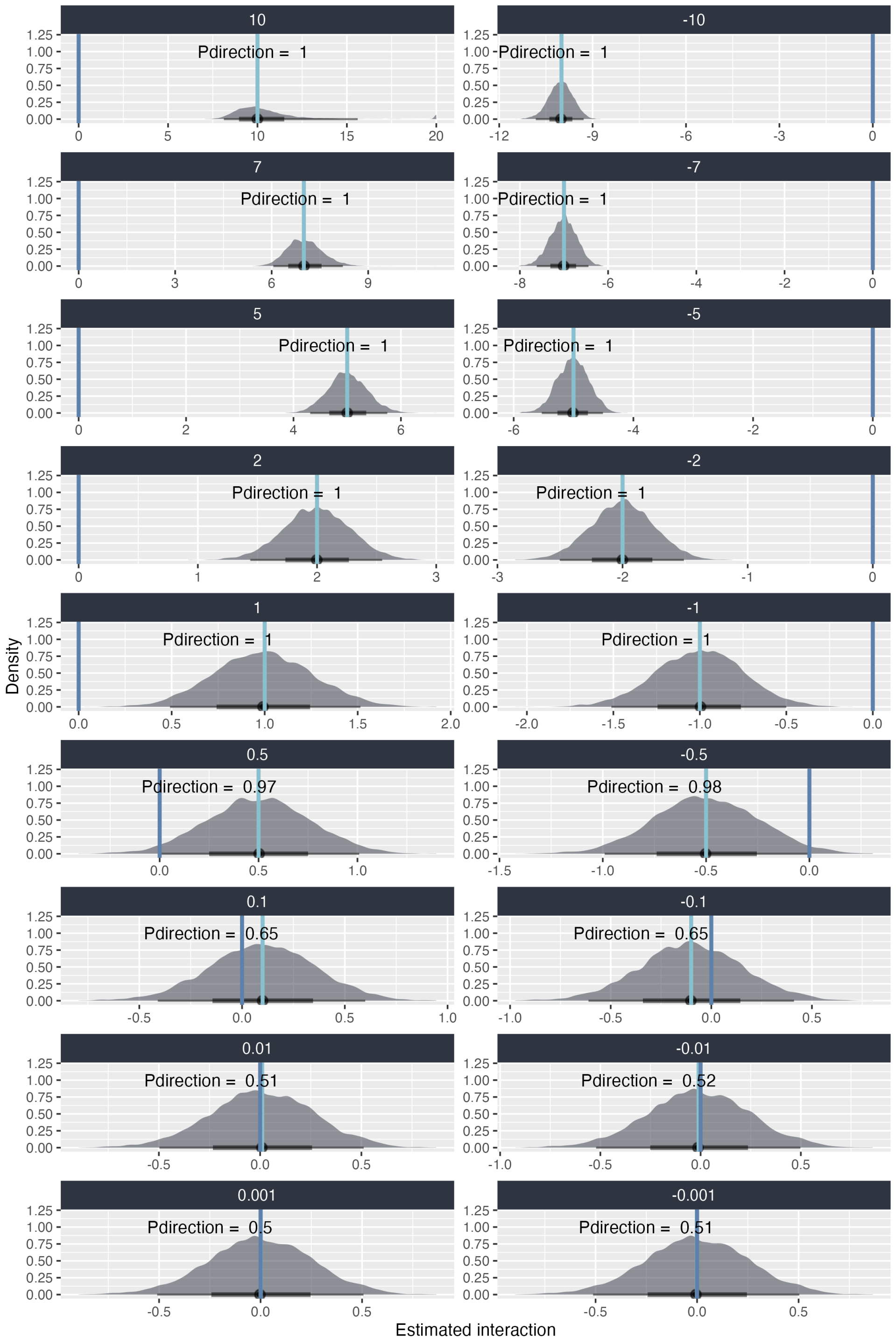

Figure 3.1: Density of error on the estimated interaction parameter using the SA model with CA as reference, for the experimental design with four replicates and different simulated values of the interaction parameter. Each panel corresponds to one simulated value, indicated on the right. Vertical light blue bars indicate the true simulated interaction value, and the probability of direction for each distribution is shown above the corresponding panel.

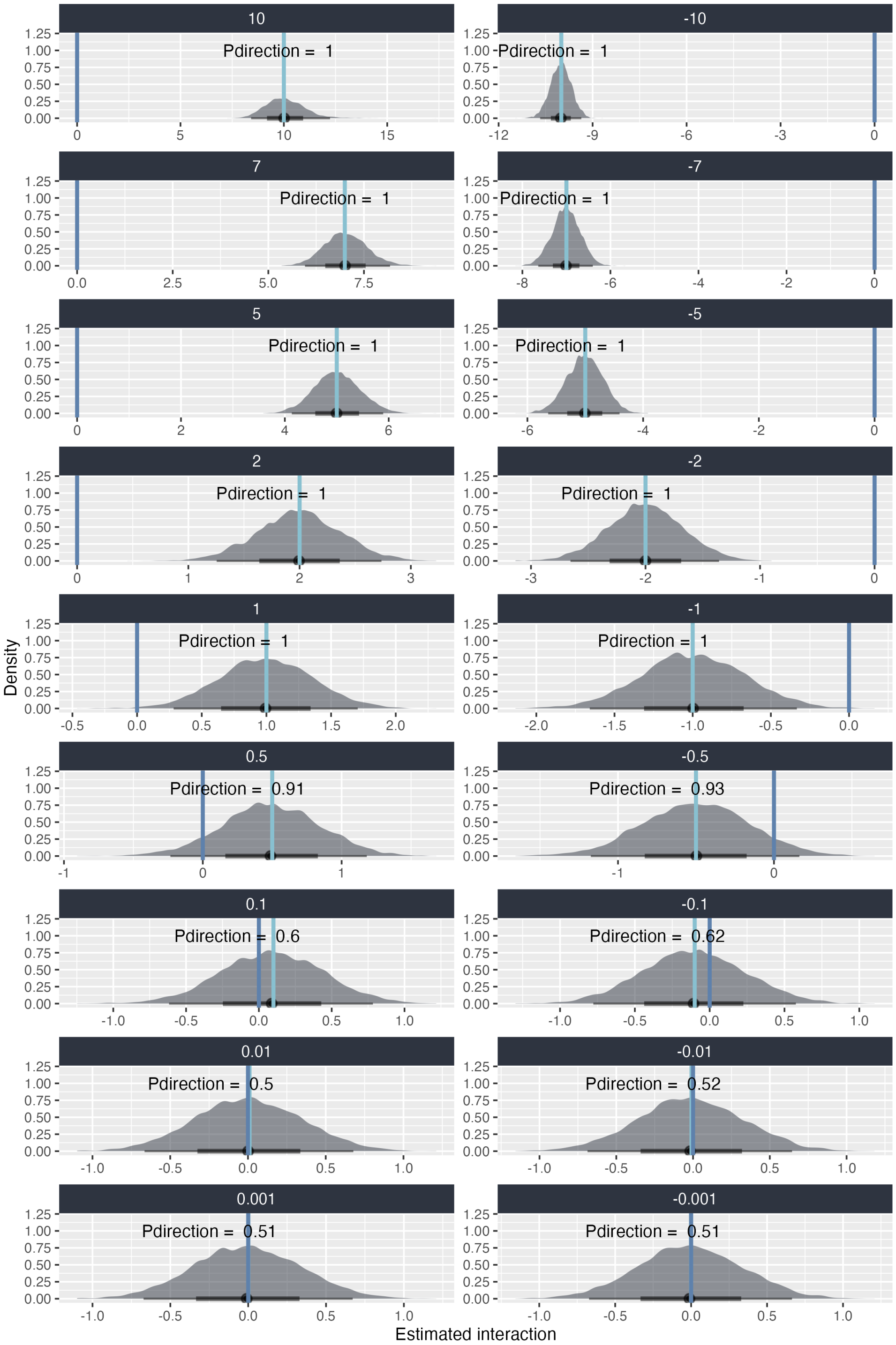

Figure 3.2: Density of error on the estimated interaction parameter using the SA model with IA as reference, for the experimental design with four replicates and different simulated values of the interaction parameter. Each panel corresponds to one simulated value, indicated on the right. Vertical light blue bars indicate the true simulated interaction value, and the probability of direction for each distribution is shown above the corresponding panel.

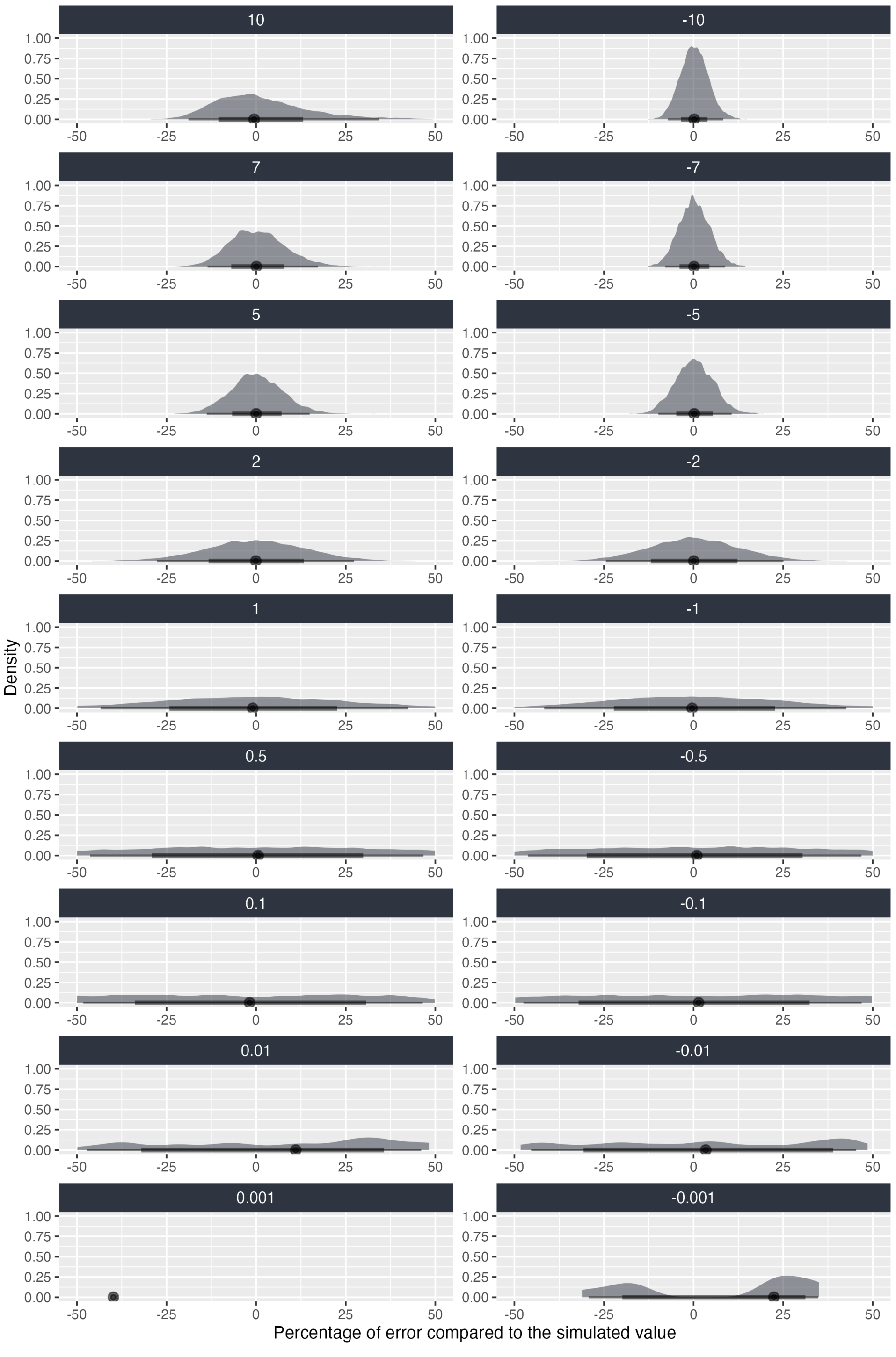

Figure 3.3: Density of percentage errors in the estimation of the interaction parameter using the SA model with CA as reference, for the experimental design with four replicates and different simulated values of the interaction parameter. Each panel corresponds to one simulated value, indicated on the right of the panels.

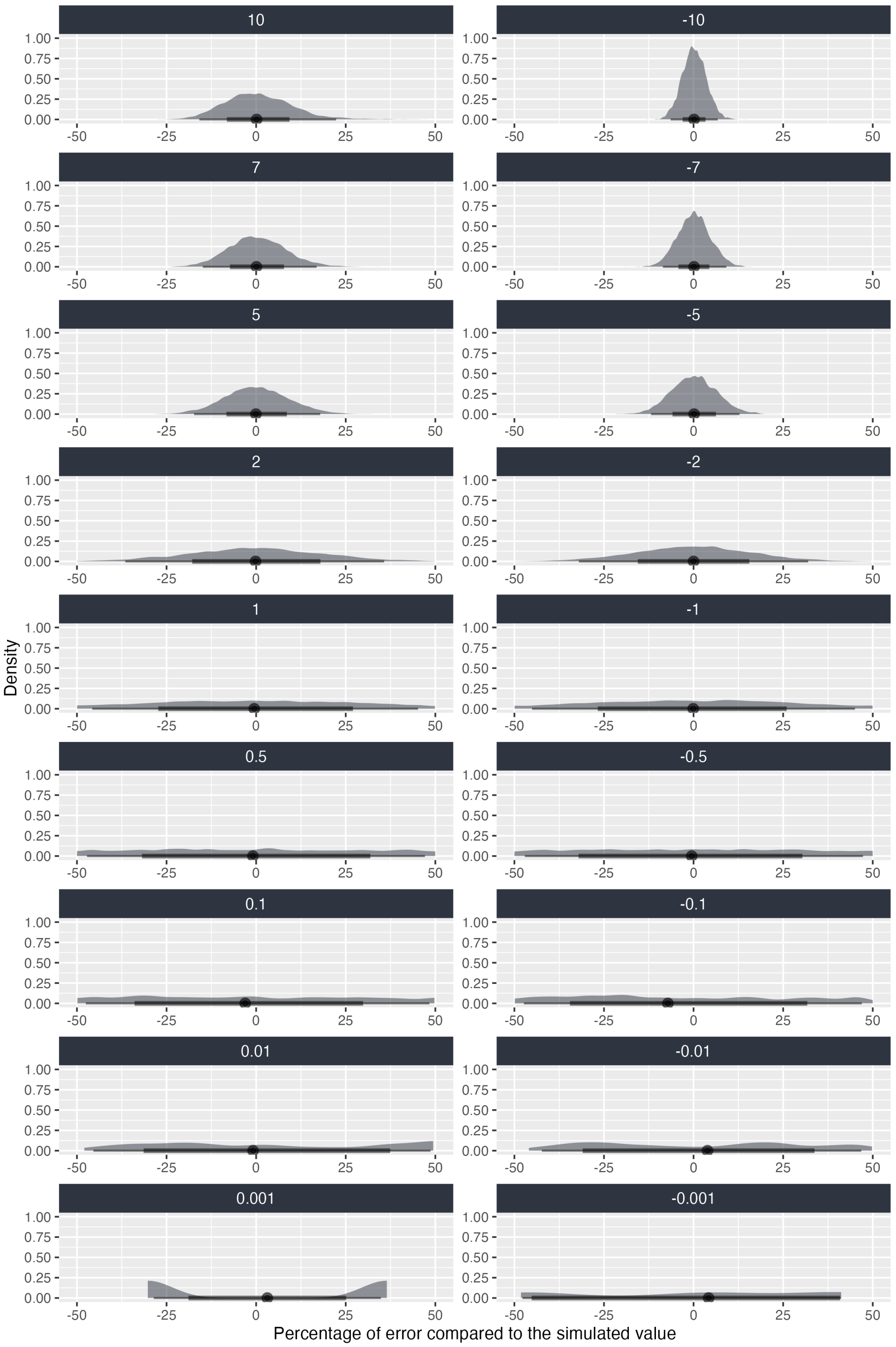

Figure 3.4: Density of percentage errors in the estimation of the interaction parameter using the SA model with IA as reference, for the experimental design with four replicates and different simulated values of the interaction parameter. Each panel corresponds to one simulated value, indicated on the right of the panels.

Table 3.1: Power, misclassification and estimation accuracy of the SA model across interaction strengths under the CA and IA reference models.

| Interaction value | Power | Misclassification rate (when estimated interaction is significant) | Median estimation relative error | >25% error in the estimated interaction value | >50% error in the estimated interaction value |
|---|---|---|---|---|---|
| **CA as reference** | | | | | |
| > 2 | 100% | 0 | < 9% | < 5-7% | < 0-0.1% |
| 1 | 97% | 0 | 20% | 32-34% | 5% |
| 0.5 | 50% | 0 | 20% | 63% | 34% |
| 0.1 | 5% | 1% | 200% | 92% | 85% |
| **IA as reference** | | | | | |
| > 2 | 100% | 0 | < 15% | < 13-19% | < 0.2-0.7% |
| 1 | 80% | 0 | 30% | 45-49% | 13-17% |
| 0.5 | 30% | 0-0.12% | 55% | 72% | 47% |
| 0.1 | 4.5% | 1-1.5% | 280% | 94.5% | 89% |

### 4 Application of the Jonker framework on our experimental data

#### 4.1 Dose-response curve prediction for each ratio

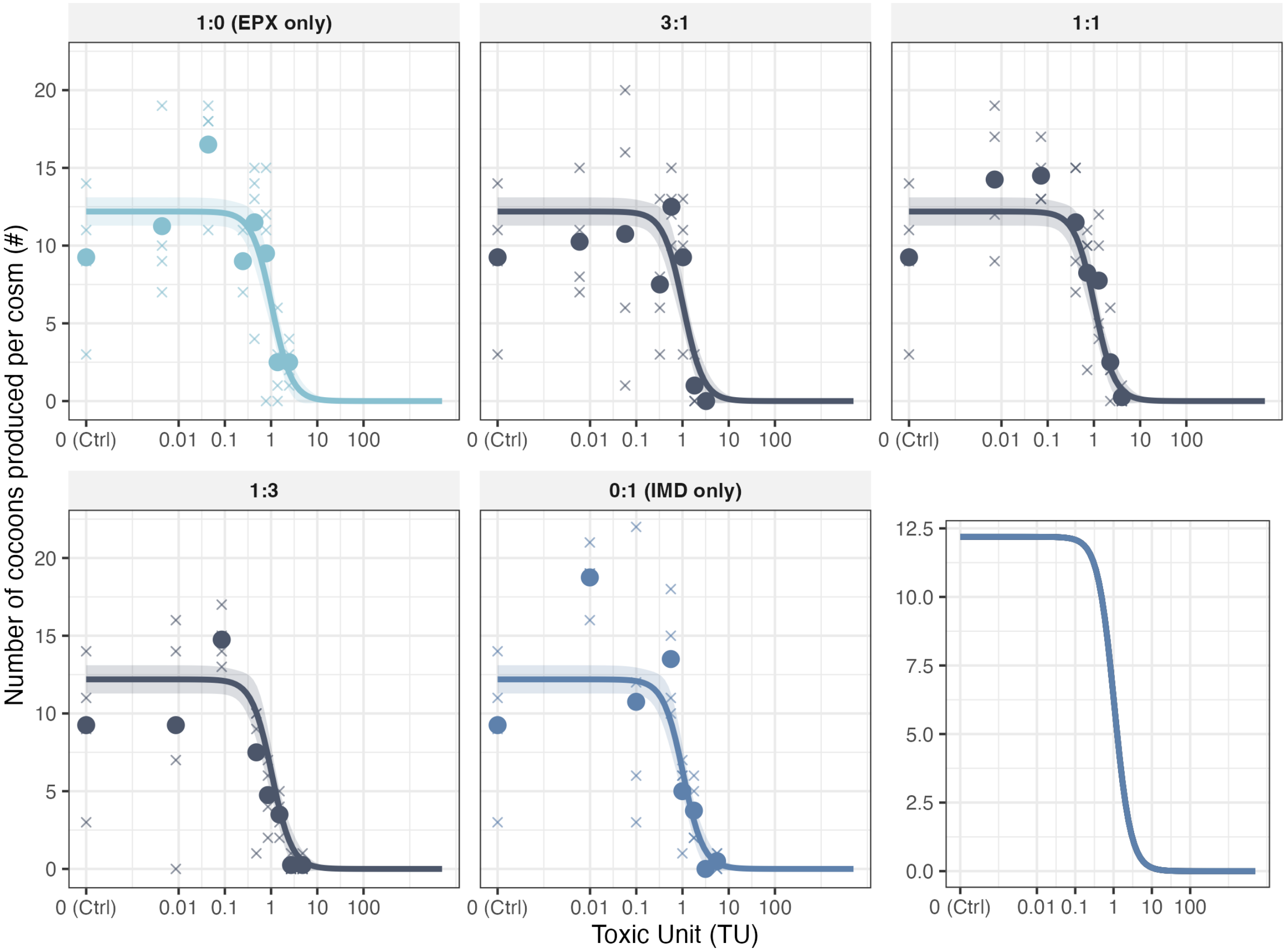

Figure 4.1: Dose-response curves obtained for each mixture ratio. The lines represent the dose-response curves obtained (with common $Y_{max}$ and $Y_{min}$ set to 0). Crosses correspond to the number of cocoons produced by one cosm and the bold points represent the mean per condition.

Table 4.1: Estimated dose-response curves for each mixture ratios. $Y_{min}$ set to 0.

| Parameter | Estimate [CI 95%] |
|---|---|
| $Y_{max}$ | 12.2 [11.3 ; 13.1] |
| Slope 1:0 (EPX only) | 2.0 [1.1 ; 2.9] |
| Slope 3:1 | 6.4 [3.5 ; 9.3] |
| Slope 1:1 | 3.0 [1.4 ; 4.5] |
| Slope 1:3 | 2.0 [1.3 ; 2.7] |
| Slope 0:1 (IMD only) | 2.65 [1.8 ; 3.5] |
| $EC_{50}$ (TU) 1:0 (EPX only) | 1.0 [0.75 ; 1.3] |
| $EC_{50}$ (TU) 3:1 | 1.2 [1.0 ; 1.4] |
| $EC_{50}$ (TU) 1:1 | 1.4 [1.0 ; 1.75] |
| $EC_{50}$ (TU) 1:3 | 0.7 [0.5 ; 0.9] |
| $EC_{50}$ (TU) 0:1 (IMD only) | 1.1 [0.9 ; 1.4] |

lacke-of-fit test with likelihood-ratio test : p value = 1.

#### 4.2 Dose-response curves for the single substance tested during experiment B

Table 4.2: Dose-response curves estimated with a different slopes.

| | Imidacloprid | Epoxiconazole |
|---|---|---|
| $Y_{min}$ | 0 (fixed) | 0 (fixed) |
| $Y_{max}$ [CI 95%] | 13.3 [11.7 ; 14.8] | 13.3 [11.7 ; 14.8] |
| $EC_{50}$ (mg/kg) | 0.27 [0.21 ; 0.33] | 198 [135 ; 262] |
| | 2.5 [1.7 ; 3.35] | 1.8 [0.9 ; 2.65] |

Log-Likelihood ratio test - Comparison of the dose-response curve models with or without common slope :

Table 4.3: ANOVA-like table

| Model | Model Df | LogLik | Df | p value |
|---|---|---|---|---|
| Different slopes | 55 | 1501.6 | 0 | |
| Common slope | 56 | 1500.8 | 1 | 0.2175 |

#### 4.3 Mixture model comparisons

Table 4.4: Estimated interaction parameters by the different Jonker interaction models.

| Model type | a | b | Log-Likelihood |
|---|---|---|---|
| CA | - | - | -395.2 |
| CA-SA | -0.005 [-1.06 ; 0.84] | - | -395.2 |
| CA-DL | 1.6 [-0.7 ; 5.5] | 0.63 [-174.4 ; 0.65] | -392.3 |
| CA-DR | -0.40 [-3.4 ; 2.3] | 0.9 [-5.1 ; 6.7] | -395.1 |
| IA | - | - | -397.5 |
| IA-SA | -1.5 [-3.0 ; -0.2] | - | -391.3 |
| IA-DL | -1.9 [-6.8 ; -0.004] | 0.6 [-298.5 ; 2.3] | -391.1 |
| IA-DR | -1.6 [-5.6 ; 2.1] | 0.3 [-8.2 ; 8.1] | -391.3 |

lacke-of-fit tests were performed on each model with log-likelihood-ratio tests (Log-likelihood anova model = -342.1) :

- Lack-of-fit test CA/Anova: D = 106.26, df = 32, p = 6.75e-10
- Lack-of-fit test CA-SA/Anova: D = 106.26, df = 31, p = 3.57e-10
- Lack-of-fit test CA-DL/Anova: D = 100.36, df = 30, p = 1.63e-09
- Lack-of-fit test CA-DR/Anova: D = 106.032, df = 30, p = 2.02e-10
- Lack-of-fit test IA/Anova: D = 110.922, df = 32 , p = 1.23e-10
- Lack-of-fit test IA-SA/Anova: D = 98.503, df = 31, p = 5.92e-09
- Lack-of-fit test IA-DL/Anova: D = 98.085 , df = 30 , p = 3.72e-09
- Lack-of-fit test IA-DR/Anova: D = 98.491 , df = 30 , p = 3.21e-09

Table 4.5: Comparison of the different models with Likelihood Ratio tests.

| | Log-Likelihood | Df | p-value |
|---|---|---|---|
| CA-SA vs. CA | -395.2 (CA : -395.2) | 1 | 0.992 |
| CA-DL vs. CA | -392.3 | 2 | 0.052 |
| CA-DR vs. CA | -395.1 | 2 | 0.892 |
| IA-SA vs. IA | -391.3 (IA : -397.5) | 1 | 0.0004* |
| IA-DL vs. IA | -391.1 | 2 | 0.0016* |
| IA-DR vs. IA | -391.3 | 2 | 0.002* |
| IA-DL vs. IA-SA | | 1 | 0.52 |
| IA-DR vs. IA-SA | | 1 | 0.91 |

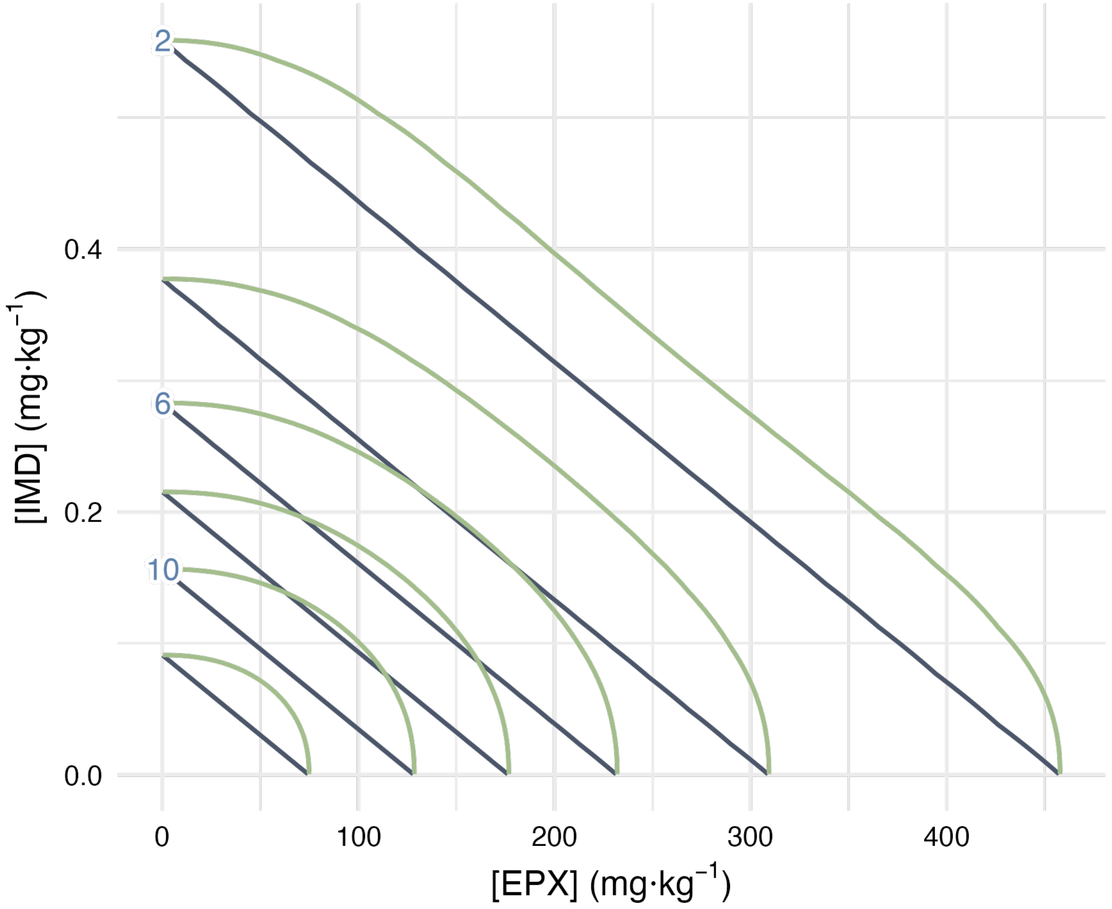

Figure 4.2: Estimated dose-response surfaces isoboles by the CA (grey) and the IA (blue) model. Blue numbers aside each isobole correspond to the level of the isobole in produced cocoons.

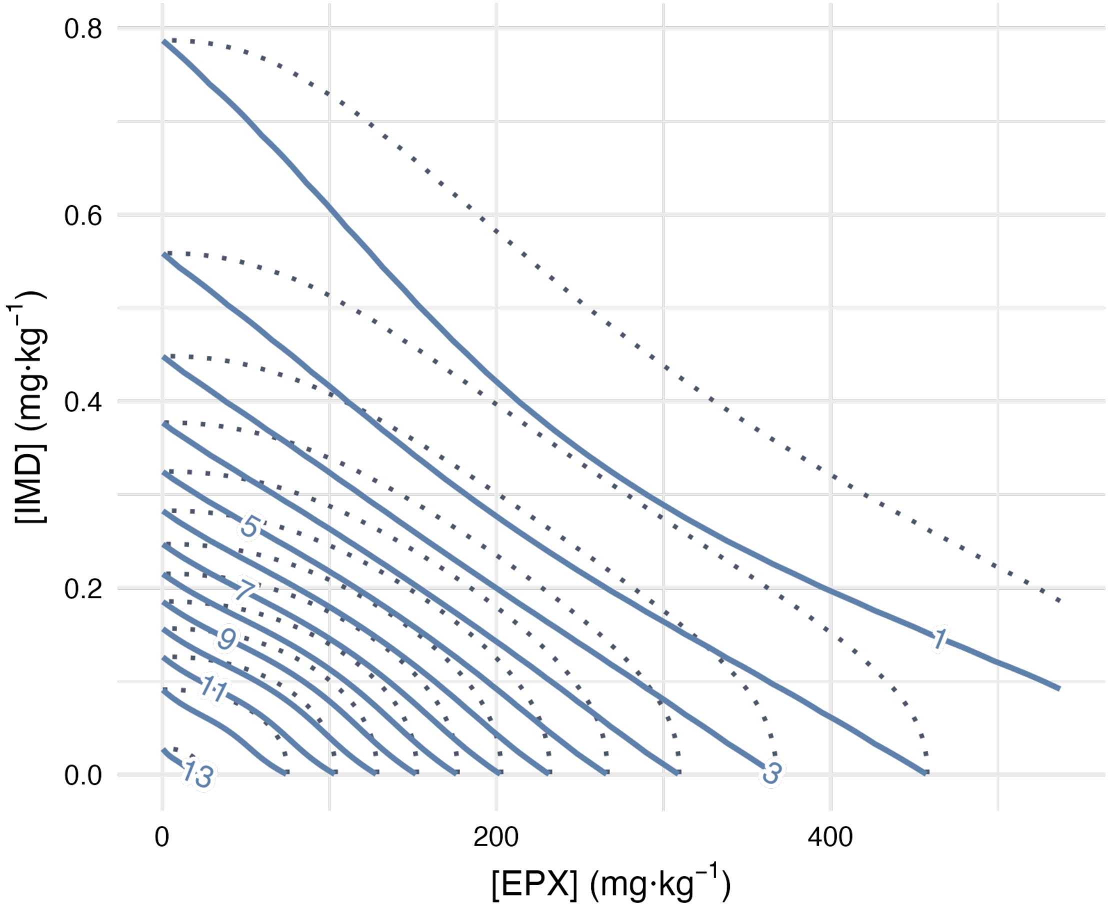

Figure 4.3: Estimated dose-response surfaces isoboles by the IA-SA (blue) model. The isoboles from the IA model are represented in dotted lines.

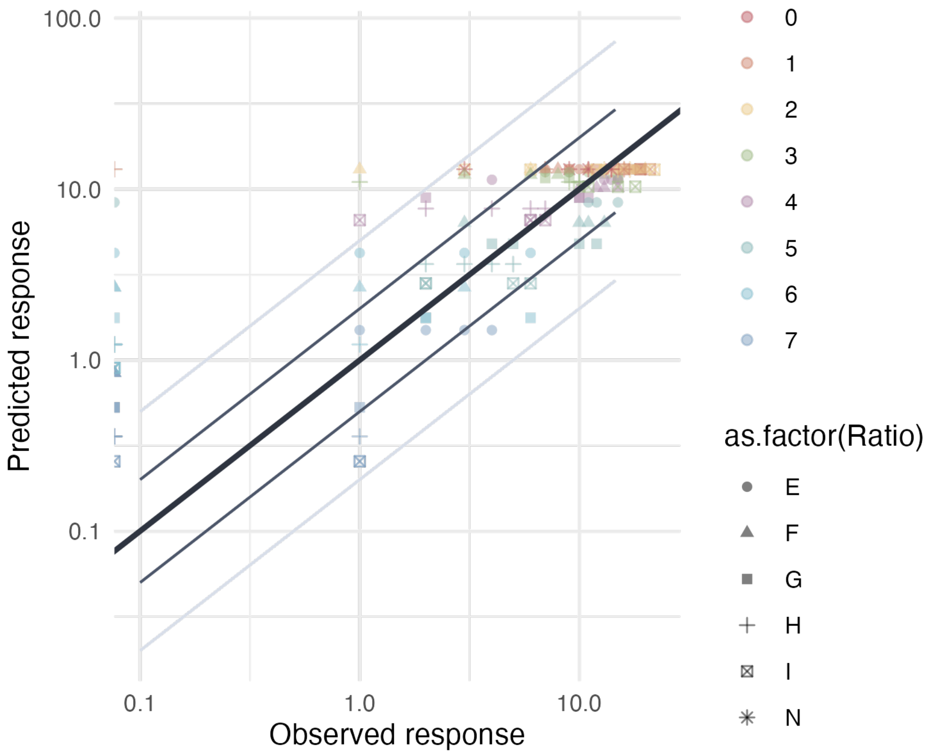

Figure 4.4: Results of the CA model fit on experimental data. Thin black and grey lines represent the 2-folds and 5-fold changes, respectively. The thicker black line represents the identity line.

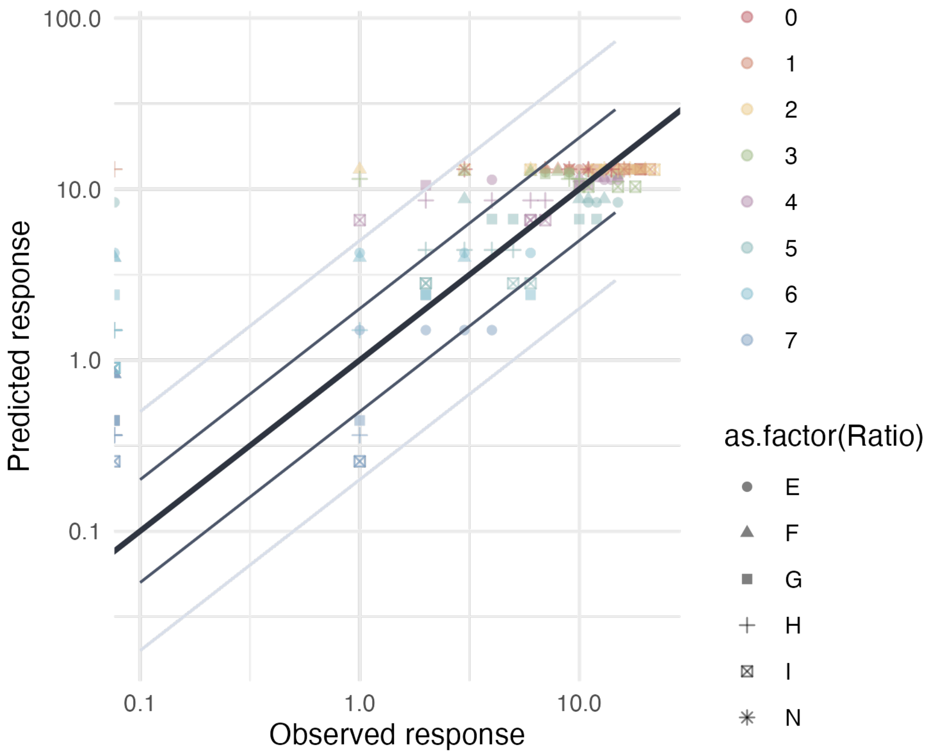

Figure 4.5: Results of the IA model fit on experimental data. Thin black and grey lines represent the 2-folds and 5-fold changes, respectively. The thicker black line represents the identity line.

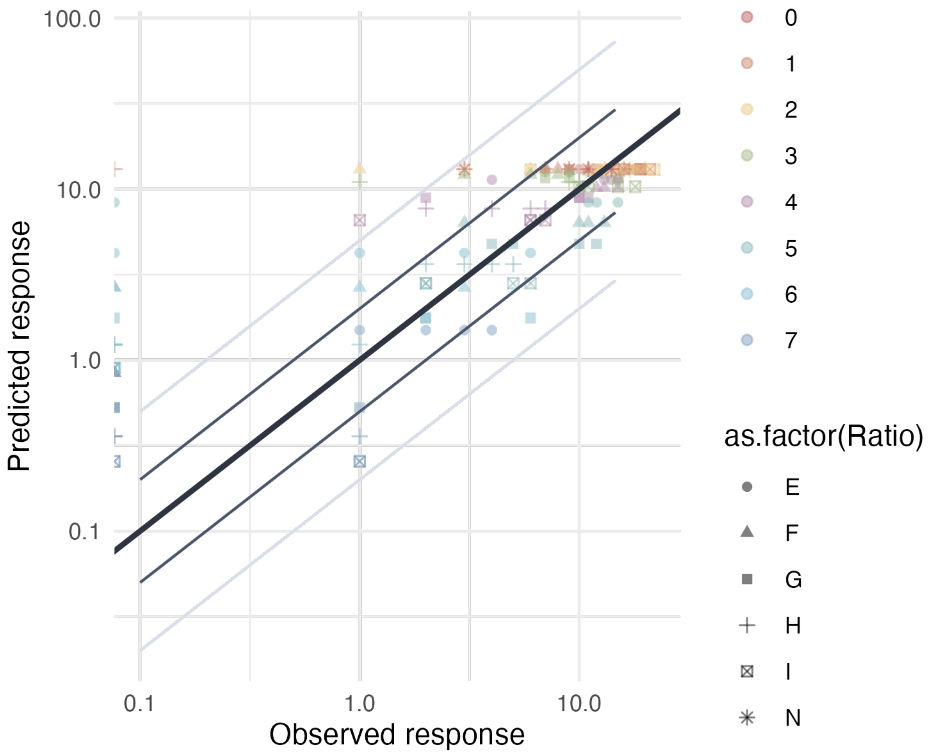

Figure 4.6: Results of the CA-SA model fit on experimental data. Thin black and grey lines represent the 2-folds and 5-fold changes, respectively. The thicker black line represents the identity line.

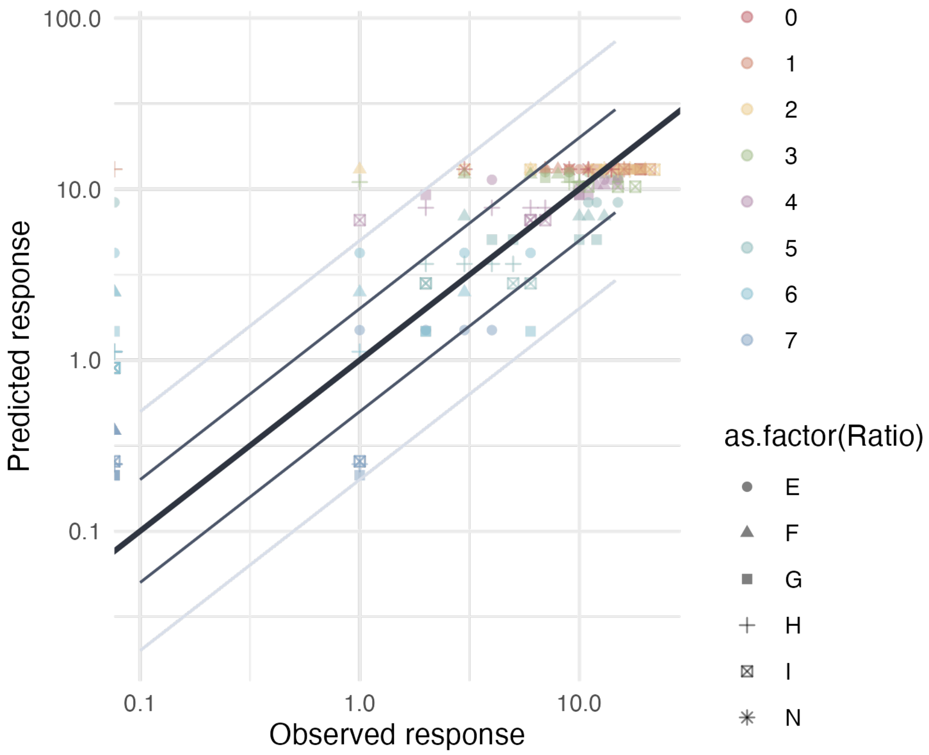

Figure 4.7: Results of the IA-SA model fit on experimental data. Thin black and grey lines represent the 2-folds and 5-fold changes, respectively. The thicker black line represents the identity line.

Predicted responses will always be strictly above zero hence the points represented on the left hedge of the plots which correspond to the cosms where no cocoon were found.
